# Renalase activates mitochondrial leak metabolism in response to cellular stress and to repair damage after injury

**DOI:** 10.1101/2025.08.19.671117

**Authors:** Xiaojia Guo, Rongmin Chen, Christine Shugrue, Fei-Fei Shang, Rolando Garcia Milian, Haley Moller, Pawel Licznerski, Tian-min Chen, Richard Kibbey, Ons M’Saad, Yuan Tian, Joerg Bewersdorf, Frank Giordano, Fred S Gorelick, Robert Safirstein, Elizabeth A. Jonas, Gary V. Desir

## Abstract

A variety of mechanisms enhance cell stress response and repair; however, the role of mitochondria in this activity is unclear. Here we show that exogenous renalase (RNLS), an intracellular flavin-dependent NADH oxidase, activates intramitochondrial RNLS activity to promote cell survival. RNLS interacts with the ATP synthase alpha and beta subunits (ATP5α and ATP5β) and opens the ATP synthase c-subunit leak channel to increase complex I and II activities and protein synthesis rate. RNLS causes a selective, sustained, time-dependent increase in cellular protein synthesis without affecting cell proliferation, whereas RNLS deletion or direct inhibition of the mitochondrial leak blocks RNLS-mediated protein synthesis. Functional analysis of newly and differentially synthesized proteins over 24 hours reveals rapid changes in one-carbon metabolism and ribosomal biogenesis pathways as early as one hour after RNLS exposure. Mitochondrial injury is more severe in the RNLS KO kidney after acute stress, related to decreased protein synthesis rate and increased mitophagy. RNLS KO mice exposed to the stress of chronic cardiac pressure overload fail to develop cardiac hypertrophy, the physiological response, and die of heart failure and cardiac rupture. These data highlight the critical role RNLS has in activating mitochondrial leak metabolism to induce selective protein synthesis and protect against acute and chronic stress.

**HIGHLIGHTS:**

- Renalase interacts with the ATP synthase alpha and beta subunits
- Renalase activates mitochondrial leak metabolism
- Renalase and leak metabolism increase complex I and II activities
- Leak metabolism increases protein synthesis rate
- Renalase protects against cell stress and organ injury

Graphical Abstract

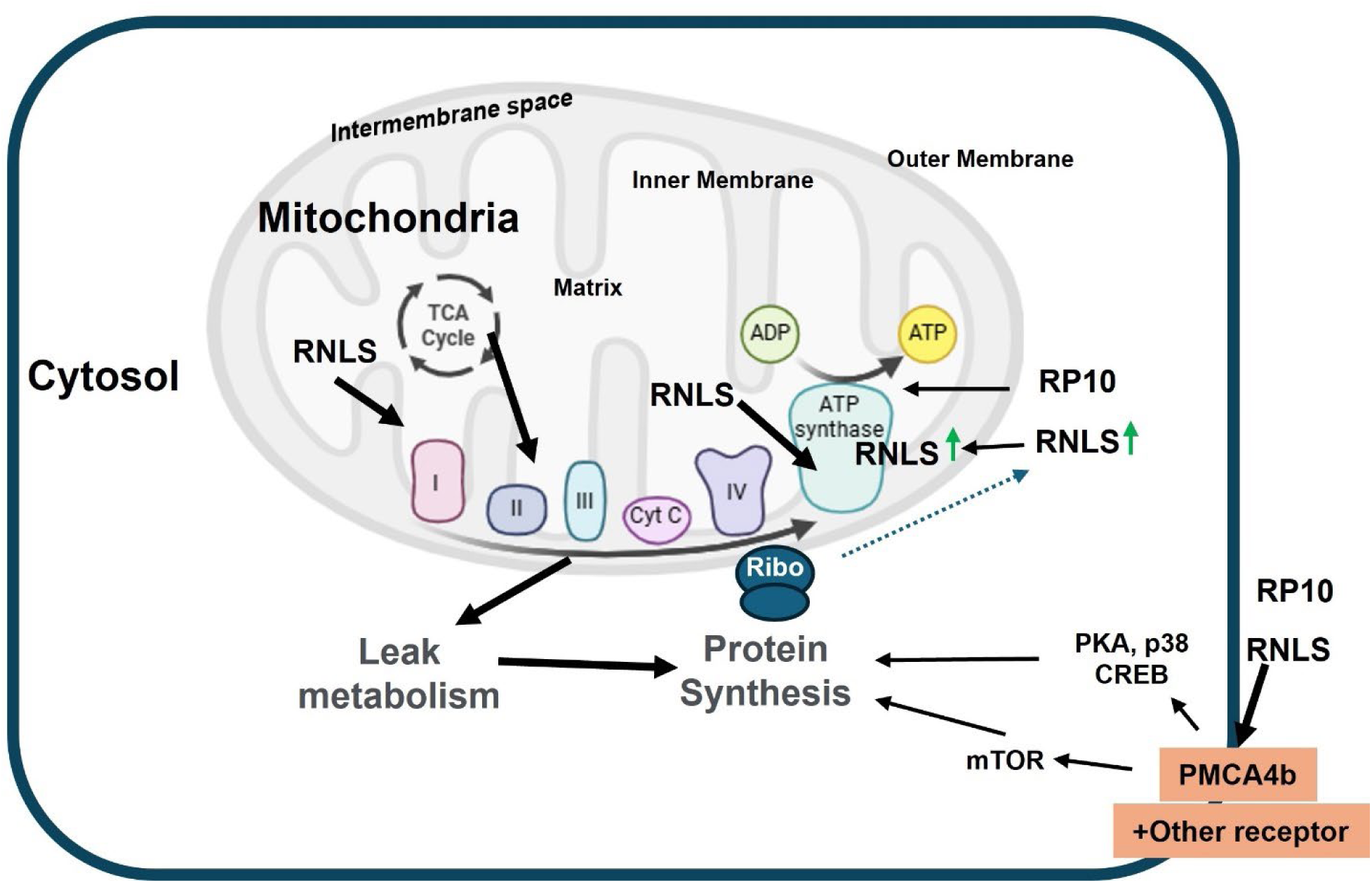

## INTRODUCTION

A range of cellular activities orchestrate responses to injury and promote cell survival ^1^. Among these are changes in mitochondrial function, enhanced protein synthesis, and upregulated clearance mechanisms. Renalase (RNLS) is a secretory protein made by renal tubule cells and other cell types ^2^. It is an ancient, highly conserved protein likely to originate from nuclear-localized plastid-like DNA fragments from cyanobacteria^3^. Its evolutionary existence and its remarkable degree of sequence conservation (up to 25% amino acid identity with bacterial proteins) suggest that it possesses fundamental function(s). RNLS contains two highly conserved, well-studied functional domains. One mediates intracellular NADH oxidation ^4^. The other interacts with ATP2b4, a ubiquitous plasma membrane calcium ATPase, and activates specific intracellular signals that promote survival in cellular and in vivo models of acute tissue injury^5,6^. A short, highly conserved peptide (aa 220 to 239 of human RNLS, designated RP220, and contained in the RNLS mimetic peptide, RP10) can function as a signaling module that replicates the cytoprotective properties and intracellular signaling pattern of recombinant human RNLS (rRNLS). rRNLS and RNLS mimetic peptides (mpRNLS) contain the RP220 sequence and enhance tissue survival after injury^7–9^. In a mouse model of renal ischemia-reperfusion injury, global genetic deletion of RNLS increased injury^10^. In wild-type mice, ischemia-reperfusion injury was associated with an ∼80% reduction in kidney RNLS expression within 24 hr. The effects of ischemia-reperfusion injury in WT mice were largely reversed by treatment with full-length recombinant human RNLS^10^. Our studies have shown roles for endogenous RNLS modulating the severity of acute renal, cardiac, and pancreatic injury ^9–11^. Similarly, we reported that RNLS can be a driver of select cancers, including melanoma and pancreatic cancer.

In one of our studies, renal injury caused by the cancer chemotherapeutic agent cisplatin was examined. Cisplatin (CP) is the most effective chemotherapy for several malignancies ^12–14^. However, renal toxicity is seen in 20-30% of individuals who receive cisplatin, which limits its therapeutic use ^3,15^. Cisplatin toxicity is associated with functional and structural damage to renal mitochondria, most notably in proximal renal tubule cells^15^. We reported that treatment with mpRNLS (RP81) containing RP220 essentially reversed cisplatin-induced renal injury and was associated with increased expression of mitochondrially encoded genes^8^. These findings suggested that RNLS could be an attractive therapeutic agent for treating acute injuries, including those caused by cisplatin, and suggested that its actions could include mitochondrial effects.

Additional preliminary observations suggested that RNLS could be essential to mitochondrial function. First, the genetic deletion of RNLS in mice is associated with a >50 % reduction in myocardial ATP and NADPH content^7,11^. Second, RNLS binds NADH/NADPH and functions as an NADH oxidase. It could regulate the availability of metabolically active NADH to affect cellular and mitochondrial processes^16–20^. Lastly, examining murine and human cell lines at low resolution suggested that RNLS immunoreactivity could be concentrated in mitochondria^10^.

Based on these findings, we hypothesized that extracellular RNLS regulates mitochondrial function. Here we show that extracellular RNLS activates a previously unrecognized survival program mediated by intracellular RNLS actions at mitochondria, characterized by a time-dependent overall increase in cellular protein synthesis rate with specific changes in proteins involved in oxidative phosphorylation, glycolysis, the tricarboxylic acid cycle (TCA), ribosomal biogenesis, and the antioxidant response. We confirm that RNLS is expressed in mitochondria and that it regulates the electron transport chain, generating a mitochondrial proton gradient and increasing cellular oxygen consumption rate. Surprisingly, RNLS also interacts with the mitochondrial ATP synthase to enhance mitochondrial inner membrane leak. We show that its action on protein synthesis is directly linked to its capacity to increase mitochondrial leak. Lastly, in two clinically relevant models, cisplatin-induced renal injury and cardiac hypertrophy in response to chronic elevation of cardiac pressure, RNLS deficiency increased injury and prevented physiological responses, including the increase in protein synthesis required for repair and compensation. These findings suggest that extracellular RNLS promotes cell survival by altering the balance between oxidative phosphorylation and aerobic glycolysis to increase cellular protein synthesis rate and preserve and renew functional mitochondria.

## RESULTS

### RNLS increases protein synthesis selectively in response to cellular stress induced by serum starvation

Injury attenuation is closely linked to increased protein synthesis ^21,22^. We have shown that mpRNLS (RP81) reduces tubular injury and oxidative stress in mouse kidneys exposed to the nephrotoxic agent, cisplatin. RNLS treatment reduced cisplatin injury and simultaneously induced the expression of mitochondrial genes encoding complex I, II, III, and V subunits ^8^. We reported a link between mitochondrial leak and protein synthesis rates ^23^. We next determined if the mitochondrial effects of RNLS relate to nascent protein synthesis by examining the effects of RNLS on protein synthesis rates in the proximal cell line TKPTS and primary cultures of renal tubule cells.

To test whether RNLS-mediated responses to stress and injury reduction were associated with changes in protein synthesis rate, we examined the effect of exogenous RNLS on serum-deprived cells. TKPTS renal tubule-derived cells were cultured in serum-free media for 24 hours ^24^, and then treated with recombinant rRNLS, an mpRNLS (RP10), or an inactive scrambled version of RP10 (scRP). The steady-state bulk protein synthesis rate was assessed after 1, 6, 12, and 24 hours by adding puromycin for 10 minutes ^25,26^. Western blot of cell lysates demonstrated that rRNLS increased the steady-state protein synthesis rate (**Figure 1A**). As shown in **Figure 1B**, compared to scRP, RP10 and rRNLS markedly increased the protein synthesis rate in TKPTS cells. The RNLS-mediated increase in protein synthesis varies over time and appears to peak at 6 hours **(Figure 1C)**. Extracellular RNLS treatment also caused increased intracellular RNLS expression at all time points (**Figure 1D**).

**Figure 1.**
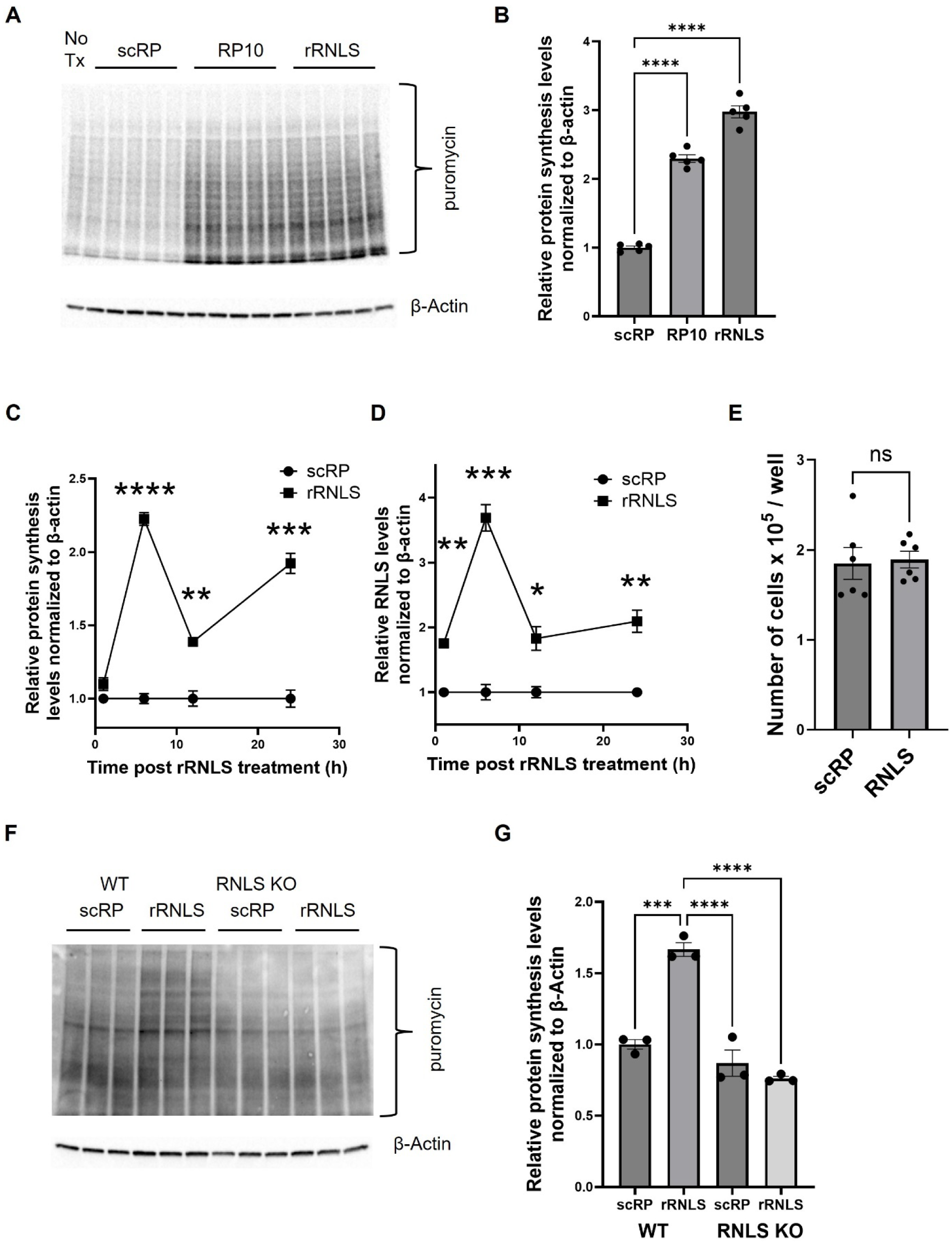
– Mitochondrial RNLS selectively upregulates protein synthesis in a time-dependent manner in response to extracellular application of rRNLS. (A) TKPTS cells were treated for 24 hours with scrambled RNLS peptide (12.5 µM; sc-RP), the biologically active RNLS peptide 10 (12.5 µM; RP10), or recombinant full-length human RNLS 1 (1.35 µM; rRNLS), and the levels of protein synthesis were measured using puromycin incorporation. A representative immunoblot of TKPTS cell lysate using anti-puromycin antibody is shown. (B) Puromycin incorporation per unit time indicates protein synthesis rate, so darker lanes reflect increased protein synthesis. Quantification of average whole lane densities is shown, n=5, ****p<0.0001by one-way ANOVA. (C-D). Quantification of immunoblots of TKPTS treated with rRNLS at the indicated times for puromycin (C) and RNLS (D). Extracellular rRNLS increases protein synthesis and intracellular RNLS level from 1 to 24 hrs (n=3, \**p*<0.05, \*\**p*<0.005, \*\*\**p*<0.001, \*\*\*\**p*<0.0001 by unpaired group t test). (E) Renalase increases cellular protein content but does not promote cell division at 24h. Viable TKPTS cell counts after 24h treatment of rRNLS or control scRP were measured. (F-G) RNLS KO Primary cultured renal cells fail to show increased protein synthesis rate upon stimulation with extracellular RNLS. Primary cultured renal cells were isolated from WT or RNLS KO kidneys and cultured in serum-free medium for 5 days, followed by a six-hour treatment with either scrambled RNLS peptide (12.5 µm; sc-RP) or recombinant full-length human RNLS 1 (1.35 µm; rRNLS), and the levels of protein synthesis were measured using puromycin incorporation. A representative immunoblot and quantification of average whole lane densities are shown (E and F, respectively), n=3, \*\*\**p*<0.001, ****p<0.0001 by one-way ANOVA.

The magnitude of RNLS’s positive impact on protein synthesis was unexpected; therefore, we determined if it was accompanied by increased cell proliferation. TKPTS cells were treated with either scRP10 or rRNLS as above, and cell counts were estimated using Trypan Blue. Cell numbers were the same at 24 hours (TKPTS population doubling time, approximately 12 hours) with and without rRNLS (**Figure 1E**), suggesting that RNLS preferentially increases cellular protein content, not cell number.

To determine if intracellular RNLS is required for the changes in protein synthesis rates, we performed the puromycin assay on primary kidney tubule cells acutely isolated from WT and RNLS KO animals. rRNLS added exogenously to the cultures increased protein synthesis rates in WT, but not RNLS KO cells (**Figure 1F, G**), suggesting that endogenously expressed RNLS acted intracellularly to increase overall protein synthesis rate.

### RNLS localizes to mitochondria and nuclei

Our previous low-resolution microscopy studies suggested that RNLS is predominantly a cytoplasmic protein ^7^. To determine if RNLS is localized to a specific subcellular compartment, we used pan-expansion microscopy (pan-ExM). This recently developed method provides exquisite subcellular details at near electron microscopy resolution using light microscopy and preserves the option to detect specific proteins by immunolabeling ^27^. When applied to mouse kidney tissue, mitochondria and cell junctions are clearly defined in proximal and distal tubules **(Figure 2A and S1A**). The glomerulus and slit diaphragms were identified morphologically by showing podocin immunoreactivity **(Figure S1B and C)** ^28^. Commercially available antibodies raised against human RNLS (rRNLS) revealed RNLS immunoreactivity is concentrated in renal tubule cell mitochondria (**Figure 2A**).

**Figure 2:**
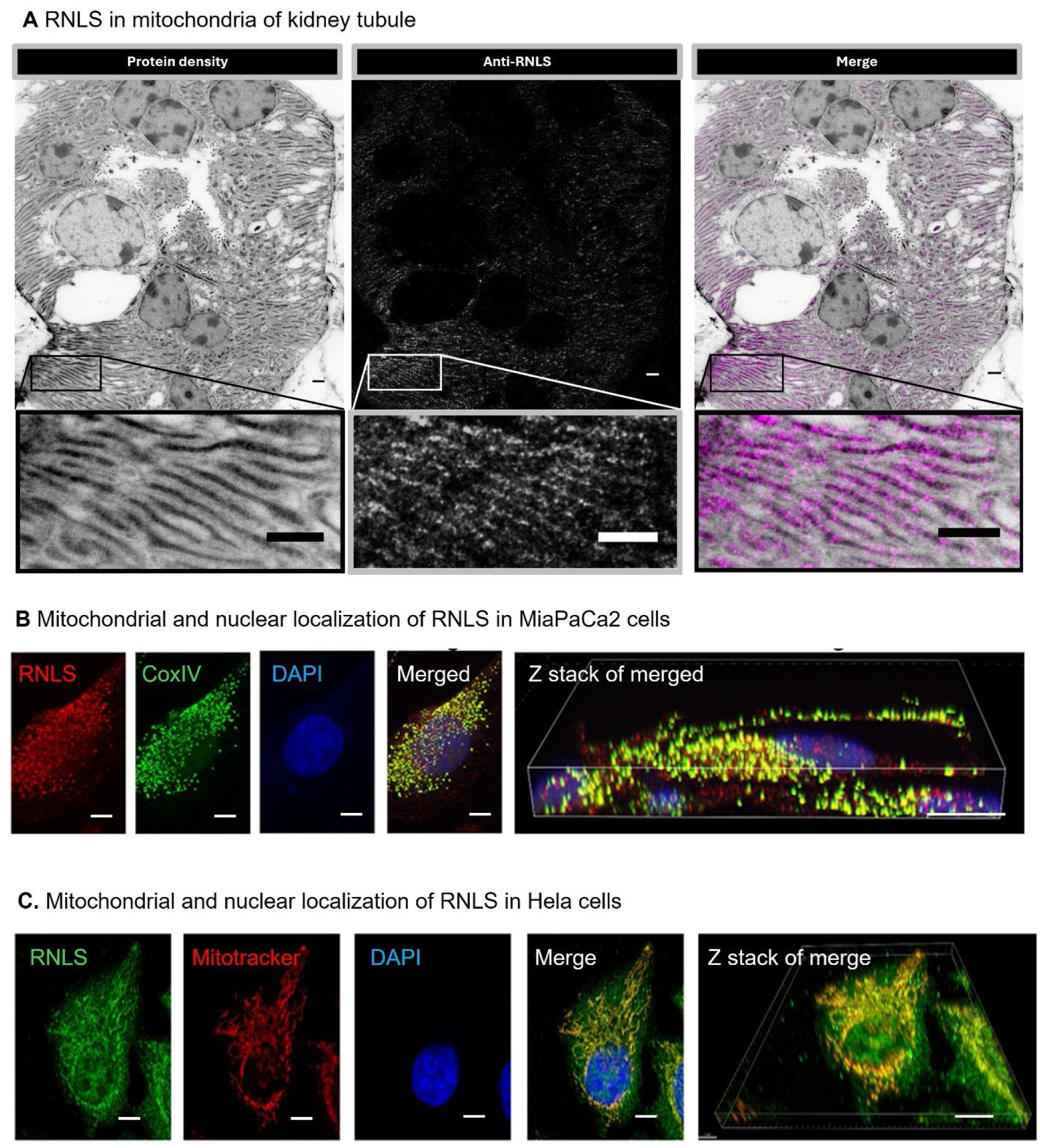
RNLS localizes to mitochondria and nuclei. (A) RNLS in the mitochondria of kidney cells. Mouse cortical proximal tubule kidney tissue revealed by Pan-expansion microscopy; protein density is displayed in a black-to-white color scale (left panel). RNLS (white in the center panel and purple pseudo colored in the right panel) is detected in mitochondria of kidney cells using a polyclonal antibody for RNLS (insets show intramitochondrial presence of antibody). Scale bar = 1 µm. (B) Mitochondrial and nuclear localization of RNLS in MiaPaCa2 cells using a monoclonal antibody against RNLS (m28) by confocal microscopy. 3-D image reconstruction shows colocalization of RNLS and CoxIV. RNLS labeling is observed primarily in mitochondria, but also in the nucleus and cytoplasm (red stain in 3-D image). Scale bar = 10 µm. (C) Mitochondrial and nuclear localization of RNLS in HeLa cells shown by confocal microscopy. RNLS and mitotracker (red) overlapped (merged). Merged images show limited RNLS immunoreactivity in nuclei, while the Z-stack image shows the predominant RNLS immunolocalization is in mitochondria. Scale bar = 10 µm.

To determine if RNLS is localized to mitochondria in another cell type, we labeled the human pancreatic cancer MiaPaCa2 cell line ^29^, which expresses high levels of RNLS. Double labeling studies using the m28 RNLS antibody and the mitochondrial marker COX IV demonstrated their co-localization, confirming the dominant mitochondrial distribution of RNLS (**Figure 2B)**. Less prominent RNLS labeling was also detected in the cytoplasm and nucleus **(Figure 2B, Z-section of Merged).** The specificity of the m28 monoclonal antibody was confirmed using kidney tissues isolated from a global RNLS KO ^11^ and epitope peptide competition **(Figure S2A and B, respectively**). The localization of RNLS to mitochondria and the nucleus was further confirmed in HELA cells overexpressing rRNLS and labeled with the m28 antibody and a mitochondrial membrane potential-sensitive dye (MitoTracker<u>^®^</u> Red CMX Ros) (**Figure 2C).** These results demonstrate that RNLS is concentrated in the mitochondria and the nucleus of several cell types. Deletion studies suggest that the RNLS N-terminal 22 amino acids are sufficient for mitochondrial targeting (**Figure S2C**). RNLS, therefore, is present primarily in mitochondria and the nucleus. Regulated “bimodal” targeting of proteins with both endoplasmic reticulum signal sequences and mitochondrial targeting signals, as seen with RNLS, has been described ^30,31^. To investigate functional roles for mitochondrial RNLS, we first considered its known ability to metabolize NADH.

### RNLS stimulates the activity of Complex I and Complex II

In vitro studies indicate that RNLS can oxidize NADH at the 2-or 6-position of the nicotinamide ring rather than at the metabolically active 4-position ^32^. RNLS and pyridoxamine-phosphate oxidase are the only enzymes known to prevent the buildup of inactive NADH substrates by converting metabolically inert 2 or 6 NADH to active 4-NAD+. Recent evidence supports a repair role for RNLS and pyridoxamine-phosphate oxidase in intact cells ^19^. Pyridoxamine-phosphate oxidase is a cytoplasmic enzyme that catalyzes the terminal, rate-limiting step in synthesizing pyridoxal 5’-phosphate, vitamin B6. Since mitochondrial ATP production through oxidative phosphorylation requires a steady supply of metabolically active NADH to support Complex I activity, loss of mitochondrial RNLS with genetic deletion could explain the dramatic decrease in cardiac ATP levels in an RNLS KO mouse^7^.

To test if RNLS could increase electron transport, we isolated mitochondria from the kidneys of RNLS KO and WT mice. We compared the mitochondrial membrane potential by measuring the uptake of the membrane potential indicator tetramethyl rhodamine ethyl ester (TMRE) in the presence of glutamate and malate to activate complex I or succinate to activate complex II without activating complex I. We found that the overall mitochondrial membrane potential without exogenous RNLS addition was not significantly reduced in RNLS KO **(Figure 3A)**. Recombinant RNLS increased the membrane potential in both WT and mitochondria isolated from RNLS KO kidney using either fuel source (**Figure 3B, C**). These data suggest that exogenous RNLS addition enhances the activities of complexes I and II. Therefore, these activities are not near their peak levels even in the presence of WT amounts of endogenous resting RNLS.

**Figure 3.**
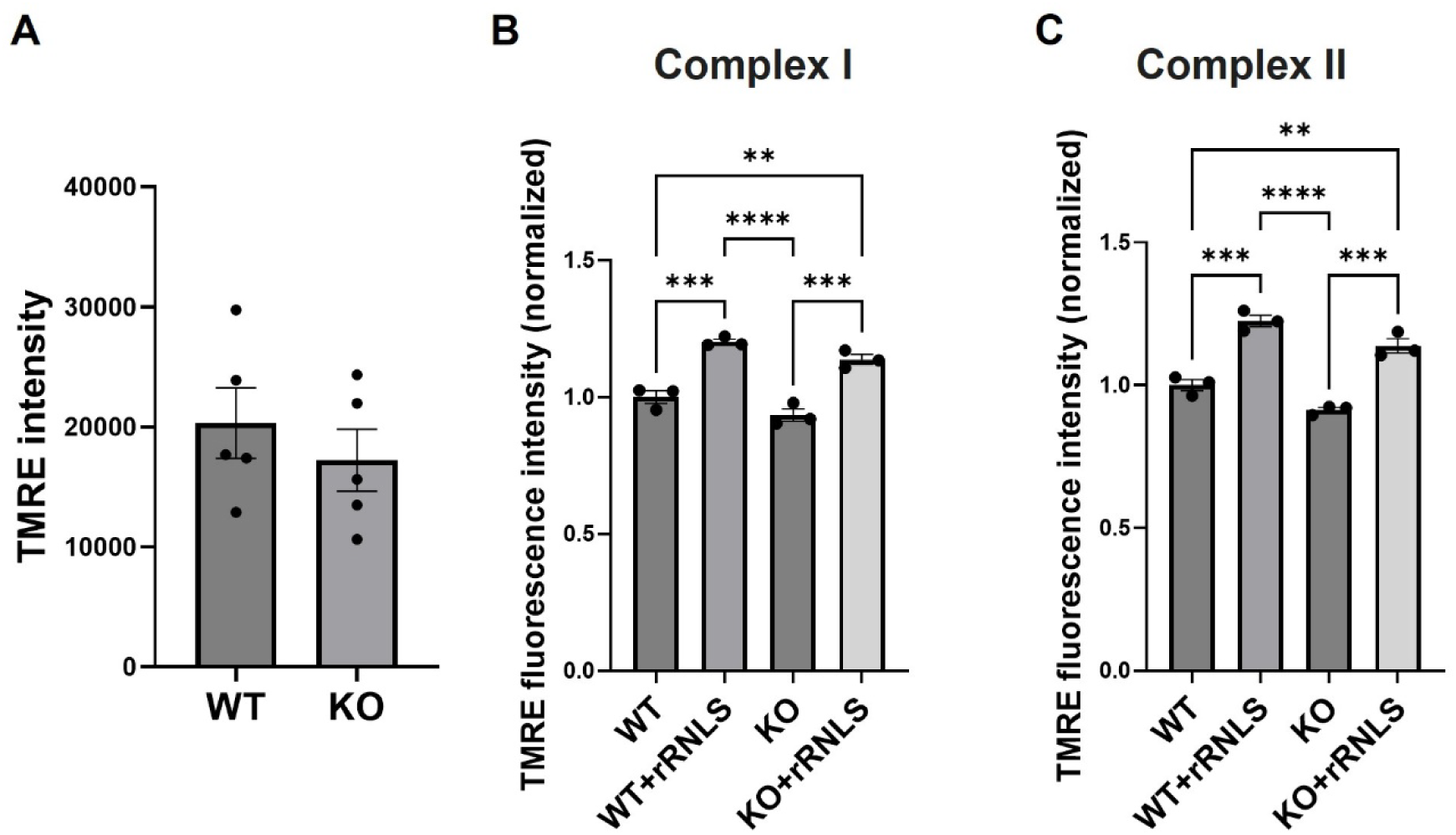
Recombinant RNLS rescues the impaired activities of mitochondrial complexes I and II in RNLS KO kidneys. (A) Membrane potential of RNLS KO kidney mitochondria trended lower than WT as measured with the membrane potential indicator tetramethylrhodamine ethyl ester (TMRE), using complex I substrates. 5 independent experiments; at least three wells for each measurement were performed. (B) rRNLS increases the activity of complexes I and II. Membrane potential (TMRE fluorescence intensity) of WT and RNLS KO mitochondria exposed to substrates specifically activating complexes I or II in the presence of rRNLS; (N=number of wells, one representative example of 5 independent experiments is shown for each complex; all data were normalized to average of WT. (**p < 0.005, ***p < 0.001. one-way ANOVA analysis of variance with a Tukey’s multiple comparisons test).

Although RNLS’s NADH oxidase activities could play a role in its effects on complex I, they are unlikely to affect complex II activity, which requires FADH2 rather than NADH. This suggests that an additional mitochondrial function could contribute to RNLS’s effect on mitochondrial membrane potential.

### RNLS opens the ATP synthase c-subunit leak channel complex and increases ATP production

RNLS may influence mitochondrial membrane potential by directly affecting the activity of ATP synthase (Complex V). To investigate this possibility and determine whether RNLS is close to specific subunits of Complex V, we used the proximity ligation assay (PLA), which detects two proteins within 40 nm of each other. Permeabilized HELA cells overexpressing rRNLS were labeled with an RNLS antibody and either an antibody against the alpha (ATP5α) or beta (ATP5β) subunits of the ATP synthase F1 catalytic domain, or against OSCP, a component of the peripheral stator. RNLS was found to be near the F1 catalytic subunit, specifically ATP5 *α* and ATP5 *β*, but not near OSCP (**Figure 4A**). These results suggest that RNLS could modulate the ATP-F1 catalytic subunit.

**Figure 4:**
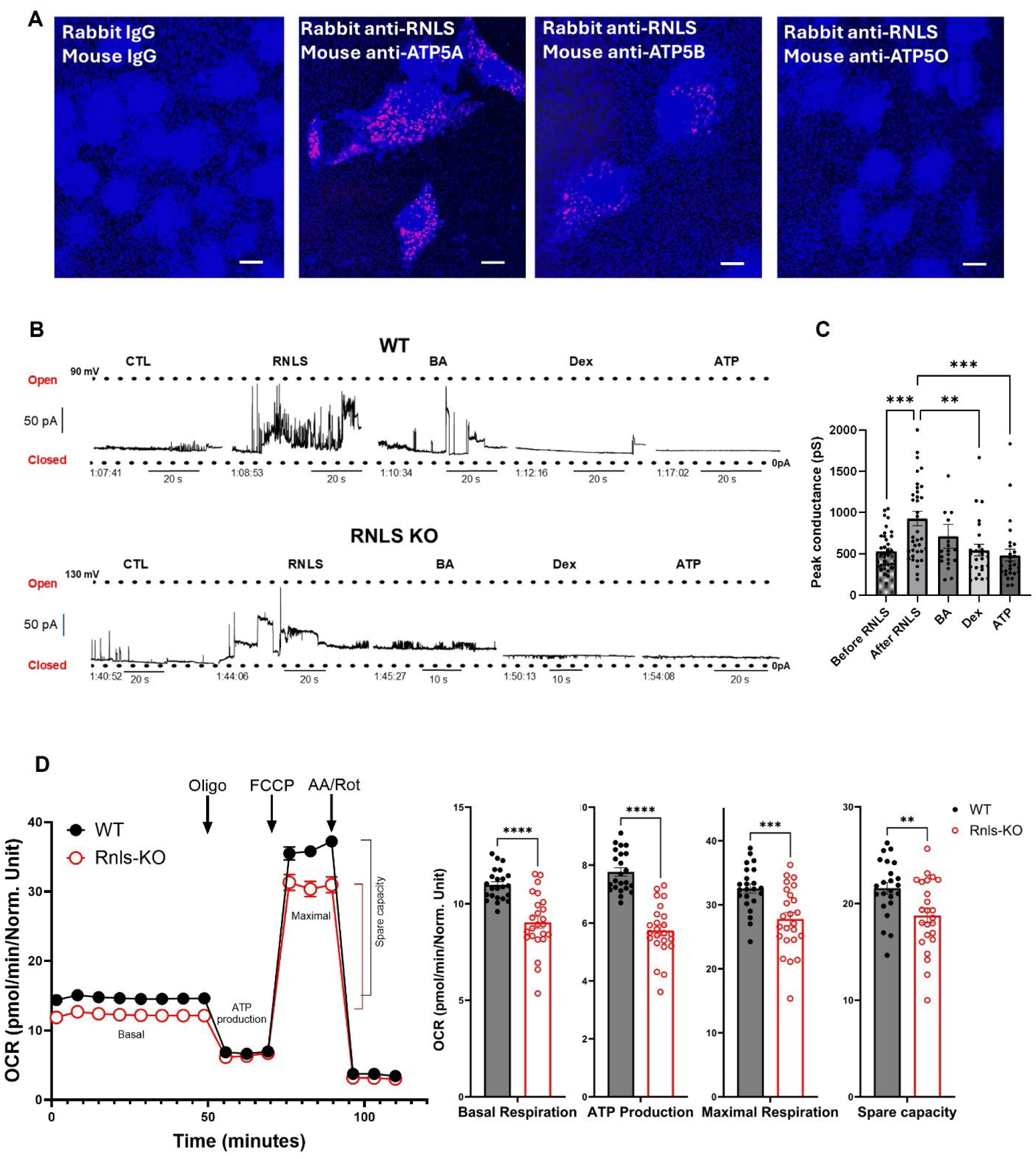
RNLS is in close proximity to the ATP synthase subunits ATP5A and ATP5B and opens the inner mitochondrial membrane leak channel. A. Proximity Ligation Assay (PLA) was performed in HeLa cells transfected with pcDNA3.1c-RNLS. RNLS is shown interacting with ATP synthase alpha and beta subunits of the stalk but not subunit OSCP of the ATP synthase stator. Scale bar = 10 µm. (B) rRNLS increased mitochondrial peak ion channel conductance in representative timed patch clamp recordings of WT or RNLS KO mitoplasts. rRNLS and inhibitors were added sequentially as indicated. (C) Quantification of all mitoplast recordings grouped together, including WT (kidney and live) and RNLS KO (one-way ANOVA; ***p=0.0010; **p=0.0064; ***p=0.0007). (D) RNLS promotes respiration and mitochondrial ATP production. Mitochondrial respiration of renal cells isolated from WT and RNLS KO mice was measured using the Seahorse XFe96 extracellular flux analyzer under the indicated conditions: oligo, oligomycin which inhibits complex V (ATP synthase); FCCP, Carbonyl cyanide-p-trifluoromethoxyphenylhydrazone, an uncoupling agent that dissipates the proton gradient; AA/Rot, Antimycin A and Rotenone which inhibits complex III and I, respectively The oxygen consumption rate (OCR) is reduced in RNLS KO renal cells compared to WT (n = 24 wells, \*\**p*<0.005, \*\*\**p*<0.001, \*\*\*\**p*<0.0001 by unpaired t test.

We have reported that the ATP synthase F1 subcomplex interacts with the ATP synthase c-subunit leak channel and inactivates it, decreasing the mitochondrial inner membrane leak and increasing the efficiency of ATP production by oxidative phosphorylation ^33,34^. Therefore, we predicted a role for RNLS in leak channel inhibition, like that of Bcl-xL binding to the ATP synthase catalytic subunit ^35,36^. In contrast, in patch clamp recordings of mitoplasts isolated from mouse kidneys or liver, exogenous recombinant full-length RNLS markedly increased the activity of the inner mitochondrial membrane leak channel within seconds of its addition to the bath (**Figure 4B, C**). Subsequent additions of dexpramipexole, bongkrekic acid, and ATP, known inhibitors of the inner membrane leak complex, rapidly closed the channel upon bath application ^34,37,38^. These data, in combination with the PLA data showing the proximity of RNLS to the ATP synthase F1 subcomplex, suggested that RNLS most likely activates, not inhibits, the ATP synthase leak channel. rRNLS opened the leak channel in both WT and RNLS KO mitochondria and was equally effective on liver and kidney mitochondria (**Figure S3**). If RNLS were already tightly bound and at full concentration, additional rRNLS might not have had as strong an effect on WT mitochondria as on the mitochondria isolated from the RNLS KO animals. However, this was not observed, suggesting that the addition of RNLS provides further activation of mitochondrial processes.

Although the above mitochondrial assays did not show a difference between KO and WT, previous findings showed decreased ATP levels in the heart **i**n RNLS KO ^7^. One result of leak channel activation is a reversal of ATP synthase to hydrolyze rather than synthesize ATP ^39^. Therefore, we determined oxygen consumption rates using the Seahorse XFe96 extracellular flux analyzer. The primary cultured renal cells isolated from the RNLS KO mouse showed significantly reduced oxygen consumption levels under basal conditions, ATP-linked respiration, and spare respiratory capacity **(Figure 4D).** Although oxygen consumption to feed the leak was not explicitly found, these data suggested that RNLS increases electron transport rates and are consistent with the data shown in **Figures 3 and 4B-C**.

### Inhibition of mitochondrial leak or RNLS signaling downregulates RNLS-mediated protein synthesis

The mitoplast patch recordings and the PLA data suggested that the direct application of rRNLS opened the ATP synthase leak channel by interacting with ATP synthase F1. Previous studies of Fragile X Syndrome had linked the ATP synthase leak channel to a metabolic phenotype in which glycolytic production of ATP is favored, and protein synthesis rates are increased ^23^. We hypothesized that this leak metabolism could also serve as a physiological signal under RNLS regulation to enhance protein synthesis rates and induce repair mechanisms. If this were the case, the inhibition of the ATP synthase leak channel should decrease the RNLS-mediated protein synthesis rates. As shown in **Figure 5A-B**, dexpramipexole and cyclosporine A, inhibitors of leak current, decreased the RNLS-mediated protein synthesis rate.

**Figure 5.**
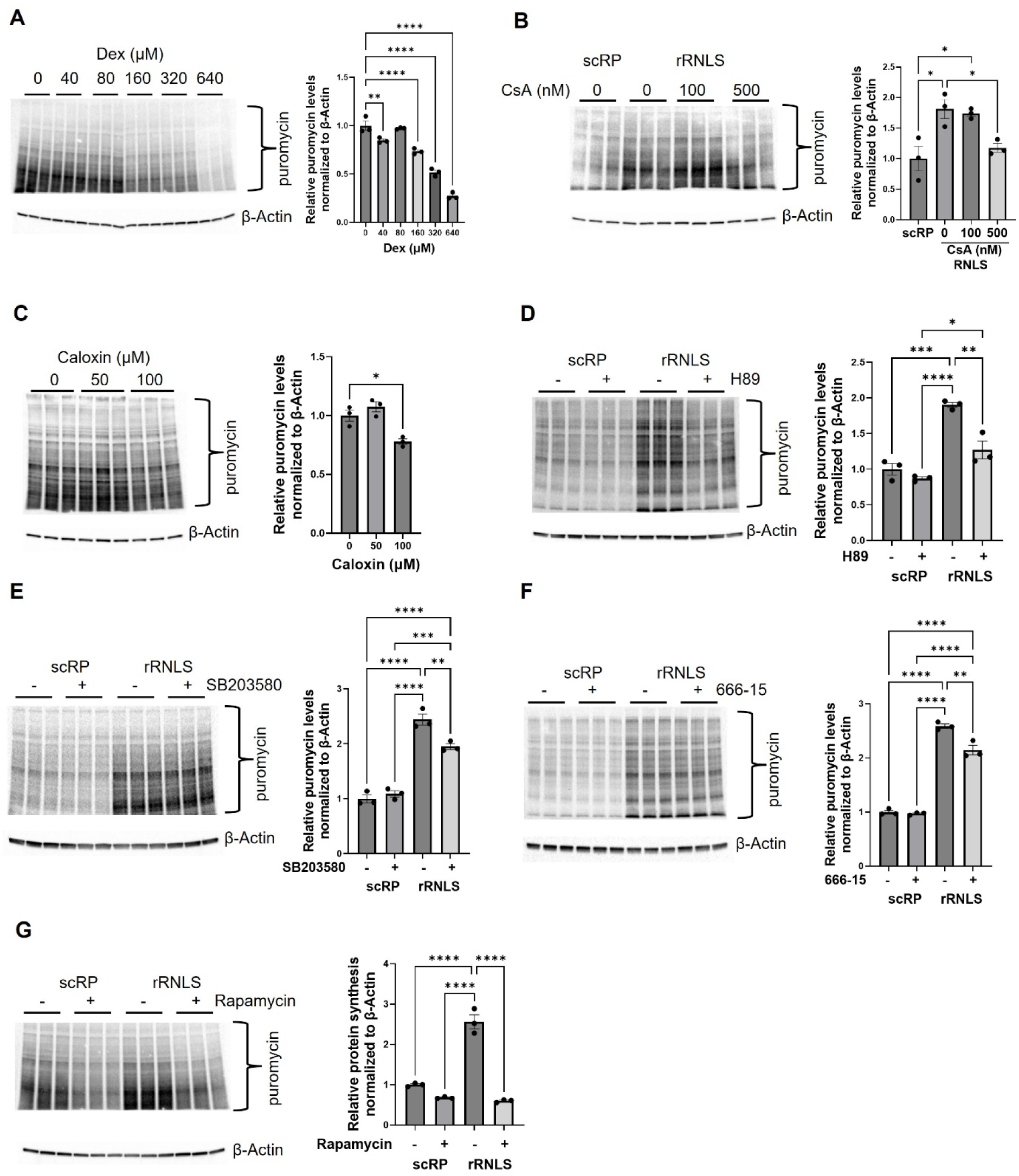
Inhibition of RNLS signaling and mitochondrial leak downregulates RNLS-mediated protein synthesis. Immunoblots (left panel) and quantification of immunoblots (right panel) of TKPTS lysate treated with inhibitors prior to the addition of rRNLS and puromycin labeling. Mitochondrial leak channel inhibitors (A) Dex (Dexpramipexole) and (B) CsA (Cyclosporin A) reduce the rRNLS-enhanced protein synthesis rate. (C) Inhibition of RNLS receptor ATP2B4 by Caloxin 1b1 reduces protein synthesis rate. (D-G) Inhibition of protein kinase A (PKA) by H89, of p38 MAP kinase by SB203580, of CREB by 666-15, and of mTOR by rapamycin reduced rRNLS-enhanced protein synthesis (n=3 cultures each, \**p*<0.05, \*\**p*<0.005, \*\*\**p*<0.001, \*\*\*\**p*<0.0001, one-way ANOVA).

Secreted RNLS is a signaling molecule that binds to its plasma membrane receptor ATP2B4 and activates cell survival pathways mediated by ERK, JAK/STAT, AKT, and p38 ^5^. Inhibition of ATPB24 by Caloxin 1b significantly decreased RNLS-mediated protein synthesis (**Figure 5C**). PKA, CREB, and p38 can be activated by extracellular RNLS and are reported to upregulate mitochondrial biogenesis ^40^. We used selective inhibitors to assess their impact on protein synthesis by decreasing PKA, p38, CREB, and mTOR activity in intact cells; the protein synthesis rate decreased by 50%, 15%, and 17% and 75% respectively **(Figures 5D-G).**

These data support the hypothesis that extracellular RNLS activates a mitochondrial leak current likely mediated by newly synthesized intracellular RNLS physically interacting with ATP5α and ATP5β. This drives elevated protein synthesis rates, perhaps through activation of PKA, CREB, p38, and mTOR downstream of ATP synthase leak channel opening.

### Proteins are differentially synthesized by RNLS

To determine if nascent protein synthesis is differentially regulated by RNLS, puromycin-containing proteins were purified from lysates obtained from control and RNLS-treated cells and subjected to mass spectrometry analysis. Of the 6,023 unique proteins identified across four time points, 797 were differentially abundant in the RNLS-treated samples compared to control samples (**Table 1 and Table S1**, two-tailed t-test False Discovery Rate (FDR) q<0.05). Principal component analysis of the differentially abundant proteins at each time point indicated minimal overlap between the experimental groups (**Figure S4A**). The non-overlapping temporal patterns suggest that extracellular RNLS induces a distinct sequence of cellular responses at different time points (**t1-24; Figure 6A-D**). We queried the STRING database (Ver 12.0) to identify protein-protein interaction networks and clusters, and to perform functional enrichment analyses for each time point ^41^. The five most enriched biological pathways involved cellular metabolism at one hour, the earliest time point (**Figure 6A and S4B**). The next five most enriched involved protein synthesis (blue lines and background Figure 6A). Components of pathways involved in carbon metabolism, including oxidative phosphorylation, TCA cycle, glycolysis, and pentose phosphate pathway, are upregulated at one hour of continuous full-length rRNLS treatment. (Table 1). Transcription and changes to DNA became more enriched at subsequent time points (6B-D). Results from the STRING database enrichment and clustering analysis of RNLS’ effect on protein synthesis again showed different enriched biological processes for each time point. KEGG (**Figure S4B**) and Reactome pathways confirmed enrichment in glycolysis, TCA cycle, and amino acid biosynthesis for cluster 1 (**Figure S4D**) and enrichment for ribosomal processes for cluster 2 (**Figure S4E**).

**Figure 6.**
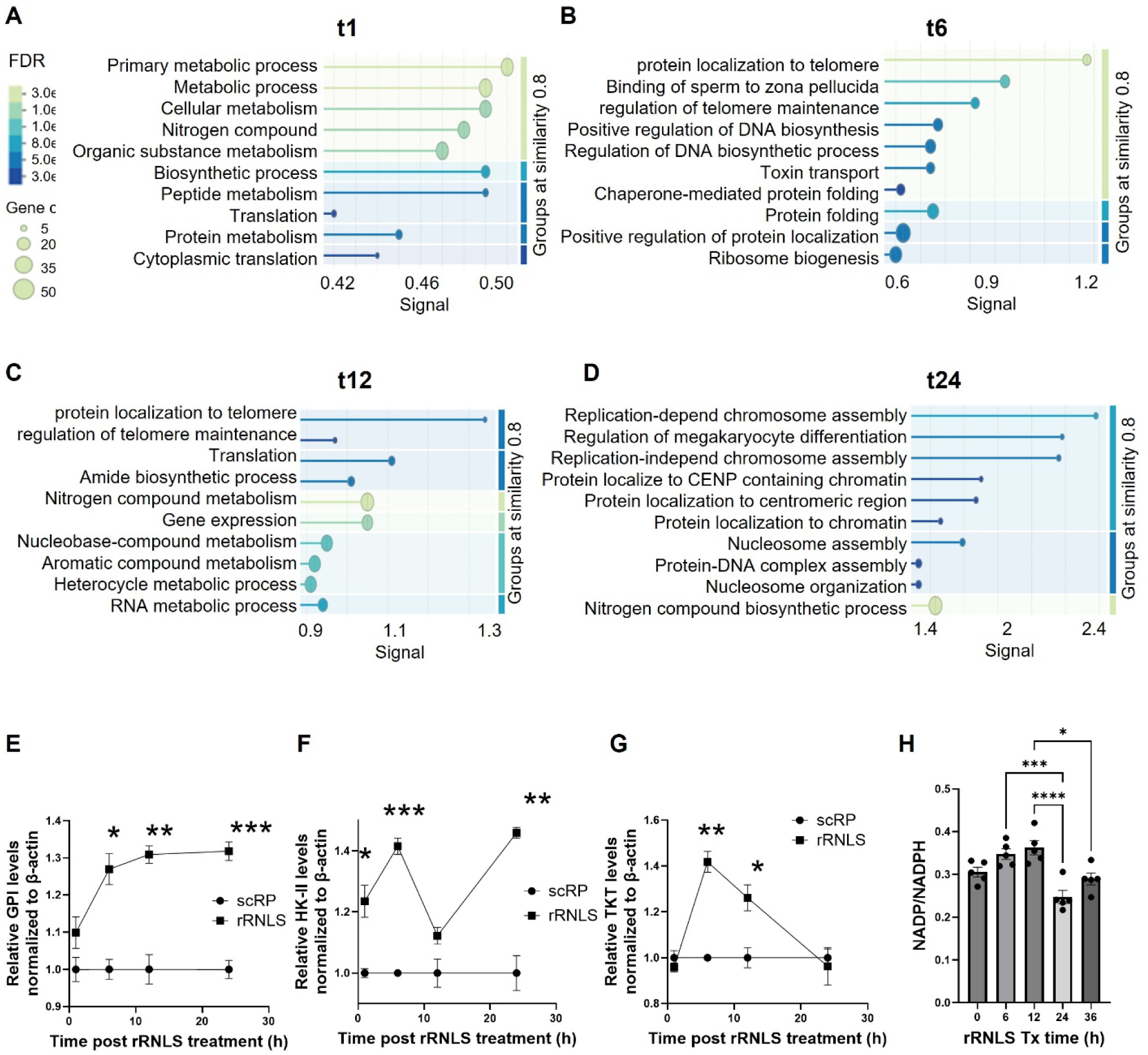
Proteins are differentially synthesized by RNLS. TKPTS cells were treated with recombinant full-length human RNLS (1.35 µm; rRNLS) for various times, followed by puromycin labeling. (A-D) The newly synthesized proteins were immunoprecipitated with an anti-puromycin antibody and subjected to mass spec analysis. Results are shown from the STRING database enrichment analysis of RNLS’ effect on protein synthesis; enriched biological processes for each time point are indicated. (E-G) Shown is the quantification of immunoblots for the indicated proteins from lysates of TKPTS cells treated with rRNLS at various times (n=3, *p<0.05, **p<0.005, **p<0.001, ****p<0.0001 by unpaired t test). Glycolytic proteins are increased by rRNLS. (H) TKPTS cells had increases in NADPH at 24 hr after exposure to extracellular rRNLS. Shown is the ratio of NADP to NADPH in TKPTS cells treated with rRNLS at various times (n=5, *p<0.05, ***p<0.001, ***p<0.0001 by one-way ANOVA).

**Table 1.** Metabolic proteins significantly changed at one hour of continuous rRNLS treatment. Data from LC/MS/MS experiments. TKPTS cells were treated with or without 1.35 μM rRNLS for 1 hour prior to puromycin treatment, followed by immunoprecipitation with anti-puromycin and processing for LC/MS/MS (n=3).

| Upregulated Protein | Metabolic Role | p-value | Log2ratio |
| --- | --- | --- | --- |
| Acads (Acyl-CoA dehydrogenase, C-2 to C-3 short chain) | Catalyzes the initial steps in the mitochondrial fatty acid β-oxidation pathway, converting short-chain acyl-CoA into enoyl-CoA. | 0.0035108 | 0.244 |
| Asl (Argininosuccinate lyase) | Participates in the urea cycle, converting argininosuccinate to arginine and fumarate, thus facilitating the removal of excess nitrogen from the body. | 0.0000001 | 16.801 |
| Atp5pb (ATP synthase peripheral stalk subunit) | Part of ATP synthase, essential for the synthesis of ATP from ADP and inorganic phosphate during oxidative phosphorylation in mitochondria. | 0.0044734 | 0.354 |
| Gsto1 (Glutathione S-transferase omega-1) | Involved in the detoxification of electrophilic compounds by conjugating them with glutathione, and also participates in the mitochondria's metabolic processes. | 0.0091297 | 0.352 |
| Hsd17b7 (17β-Hydroxysteroid dehydrogenase type 7) | Involved in the biosynthesis of cholesterol and other sterols, as well as the conversion of steroids into their active forms. | 0.0000107 | 1.838 |
| Lpcat3 (Lysophosphatidylcholine acyltransferase 3) | Catalyzing the conversion of lysophosphatidylcholine to phosphatidylcholine in phospholipid metabolism. | 0.0000045 | 15.617 |
| Mccc1 (Methylcrotonoyl-CoA carboxylase 1) | Catalyzes the carboxylation of methylcrotonoyl-CoA to 3-methylglutaconyl-CoA in the leucine degradation pathway which is essential for energy production from protein metabolism. | 0.0021860 | 1.925 |
| Mdh1 (Malate dehydrogenase 1) | Catalyzes the reversible conversion of malate to oxaloacetate, a critical reaction in the tricarboxylic acid (TCA) cycle, also known as the citric acid cycle. | 0.0012016 | 0.499 |
| Mrps25 (28S ribosomal protein S25, mitochondrial) | Part of the mitochondrial ribosome, essential for mitochondrial protein synthesis. | 0.0065786 | 0.385 |
| SLC25A5 (Solute Carrier Family 25 Member 5) | SLC25A5 encodes the ADP/ATP translocase 2 (also known as ANT2 or adenine nucleotide translocator 2) which is located in the mitochondrial inner membrane. | 0.0002763 | 2.886 |

| Downregulated Protein | Metabolic Role | p-value | Log2ratio |
| --- | --- | --- | --- |
| Aco2 (Aconitase 2, mitochondrial) | Catalyzes the isomerization of citrate to isocitrate via cis-aconitate in the TCA cycle. | 0.0005950 | -0.251 |
| Gls (Glutaminase) | Converts glutamine to glutamate providing an important source of energy and biosynthetic precursors for rapidly proliferating cells. | 0.0047821 | -0.406 |
| Acox1 (Acyl-CoA oxidase 1, palmitoyl) | Catalyzes the first step in the peroxisomal β-oxidation pathway of fatty acid metabolism, specifically acting on very long-chain fatty acids. | 0.0010505 | -0.880 |
| Gapdh (Glyceraldehyde 3-phosphate dehydrogenase) | Catalyzes the sixth step of glycolysis and thus plays an important role in energy production through the breakdown of glucose to pyruvate. | 0.0000010 | -21.188 |

We examined the time-dependent translation of selected enzymes in glycolysis and the pentose phosphate pathway using Western blot analysis (**Figure 6E-G**). Glucose-6-phosphate isomerase and Hexokinase II, which control the first and second steps of glycolysis, increased within one hour. Hexokinase may translocate to mitochondria at the same time as RNLS (compare Figure 1D to Figure 6F) and regulate protein synthesis rates (compare Figure 1C to 6F). In contrast, the increase in transketolase, a key enzyme in the pentose phosphate pathway, was transient and occurred later at 6 hours. Increased flux through the pentose phosphate pathway would raise NADPH levels, enhance antioxidant capacity, and produce ribose-5-phosphate and sugar-phosphate precursors, which are building blocks of nucleic acids. These data suggest that extracellular RNLS induces changes in intracellular RNLS levels that regulate cellular metabolism based on the rate of RNLS translation, promoting time-dependent and coordinated cellular events.

We then analyzed NADP+ and NADPH levels because they are vital molecules involved in key cellular processes, including maintaining redox balance and biosynthesis of many biomolecules like lipids, nucleotides, and amino acids^42^. As shown in Figure 6H, rRNLS gradually increased the NADP+/NADPH ratio over 12 hours, with a decrease observed at 24 and 36 hours of treatment.

### RNLS deficiency changes mitochondrial morphology and increases mitochondrial injury and mitophagy in cisplatin-induced acute kidney injury

Acute kidney injury (AKI) is characterized by mitochondrial dysfunction ^43^. The observation that RNLS deletion is associated with more severe kidney injury in cisplatin-AKI **(increased creatinine and BUN; Figure 7A)** led us to examine renal mitochondrial morphology, biogenesis, and mitophagy in the RNLS KO animals with AKI compared to WT animals. We hypothesized that an increase in protein synthesis rate in cisplatin-AKI would enhance the repair of injured mitochondria, mitigating AKI-induced injury. To evaluate injury, kidney tissue from WT and RNLS KO mice at baseline and 4 days after cisplatin-induced AKI was processed for transmission electron microscopy. Examples of the proximal tubule (renal S3 region) are shown in **Figure 7B**. Quantification of mitochondrial morphology in proximal tubule duct cells revealed signatures of repair in the WT at 4 days after injury, including a marked increase in the number of small electron-dense mitochondria consistent with mitochondrial biogenesis and leak metabolism (**Figure 7B-D**) ^23^. Prominently seen were increased lipid stores and an increased association between mitochondria and ER structures (**Figure 7E**), consistent with mitochondrially regulated high protein and lipid biosynthesis rates in the cytoplasm.

**Figure 7.**
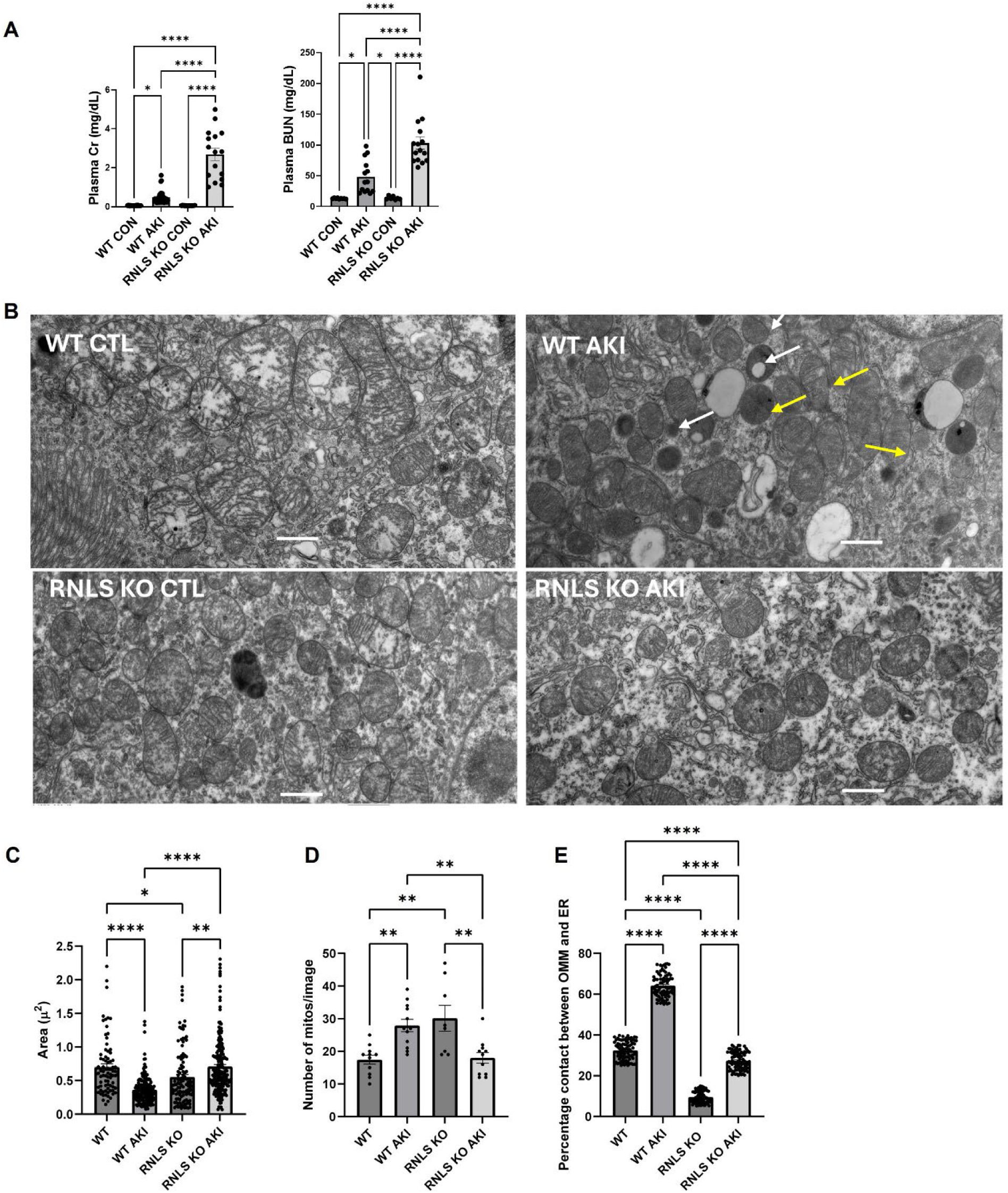
RNLS deficiency alters mitochondrial morphology and augments mitochondrial injury and mitophagy after cisplatin-induced acute kidney injury. (A) Cisplatin-induced AKI is augmented in RNLS KO as compared to WT animals, as shown by elevated plasma creatinine and BUN levels (n=16, *p<0.05, ****p<0.0001 by one-way ANOVA). (B) Morphological recovery changes of WT kidney cells after cisplatin-induced injury are not present in RNLS KO kidney. Representative electron micrographs of the S3 segment of the proximal tubule in the kidneys of WT control, WT AKI, RNLS KO control, and RNLS KO AKI at 4 days after saline or cisplatin administration. White arrows indicate lipid droplets. Yellow arrows indicate rough ER. Scale bar=1 µm. (C-E) Quantification of mitochondrial morphological changes in Panel B as analyzed for mitochondrial area (left panel), number of mitochondria (middle panel), and % contact between the outer membrane of the mitochondria (OMM) and the endoplasmic reticulum (ER) (right panel). (11-23 images of each condition; *p<0.05, **p<0.005, ***p<0.001, ****p<0.0001, one way ANOVA).

In contrast to the WT, changes consistent with repair were absent in electron micrographs performed on RNLS KO after cisplatin-induced AKI. Darkening of the mitochondrial matrix failed to occur, and the association of ER and mitochondria did not increase beyond the WT control level (**Figure 7B-E**). Instead of an increased number of mitochondria, there was a marked decrease in mitochondrial number compared to baseline, consistent with increased mitophagy and reduced biogenesis.

As shown **in Figure 8**, PGC1α, HK-II, and OPA-1 protein levels were markedly decreased in RNLS KO AKI compared to WT AKI kidney, suggesting decreased mitochondrial biogenesis, reduced glycolytic flux, and impaired mitochondrial fusion (**Figure 8A-E**). On the other hand, the mitochondrial fission protein, Drp1, phosphorylated AMPKα (Thr172), HO-1, LC3-I, LC3-II, Parkin showed significantly increased expression in RNLS KO with AKI, suggesting increased mitochondrial stress and autophagy/mitophagy **(Figure 8)**. These data indicate that RNLS deletion increases mitophagy and decreases mitochondrial repair in cisplatin-induced AKI.

**Figure 8.**
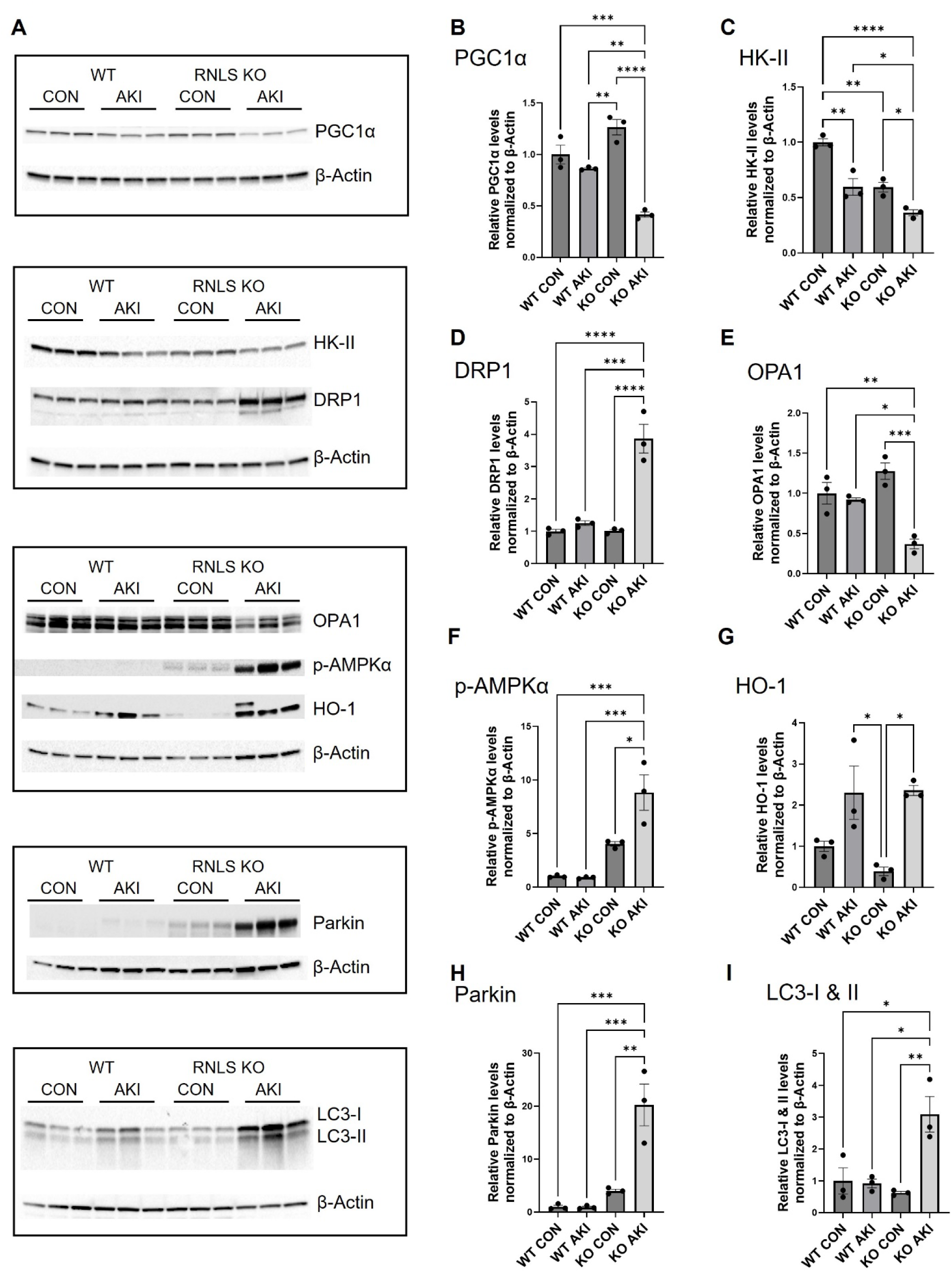
RNLS deletion augments mitophagy and attenuates mitochondrial biogenesis. Shown are immunoblots (A) and quantification of master proteins for mitochondrial biogenesis PGC1α (B), glycolytic enzyme HK-II (C), mitochondrial fusion protein OPA1 (D), cellular energy sensor, phospho-AMPKα (E), oxidative damage marker heme-oxygenase-1 (HO-1) (F), mitophagy markers Parkin (F) and LC3-I/II (G). Kidney lysates of WT and RNLS KO mice were taken 4 days after saline (control) or cisplatin (AKI) treatment (one-way ANOVA, n = 3 each, \**p*<0.05, \*\**p*<0.005, \*\*\**p*<0.001, \*\*\**p*<0.0001. β-actin was used as a loading control.

### RNLS deficiency impairs physiological cardiac hypertrophy in response to pressure overload

To determine if RNLS could modulate a different cell-stress response, we examined its role in adult terminally differentiated cardiac myocytes exposed to chronic hemodynamic overload. Such stress is induced by exercise or hypertension and results in an increased myocyte size that requires increased protein synthesis ^44,45^. Different systems and molecules have been implicated in regulating cardiac hypertrophic response, but a role for RNLS has not been examined ^44^. We hypothesized that RNLS might promote the development of cardiac hypertrophy as a physiological response to chronic pressure overload.

Cardiac pressure overload was induced in wild-type and RNLS KO mice by surgically constricting the transverse aorta (TAC) as described ^46^ (**Figure 9A**). In RNLS KO mice, we observed significant left ventricular dysfunction by 3 weeks and increased mortality from heart failure and left ventricular wall rupture (**Figure 9B-F**). There was marked cardiomyocyte loss and significant inflammation in KO hearts by 1-week post-TAC, with areas of transmural myocyte loss and severe left ventricular wall thinning by 4 weeks **(Figure 9G-H)**. These changes were not observed in wild-type TAC mice. These results suggest that RNLS is required to develop cardiac hypertrophy in response to chronic pressure overload.

**Figure 9.**
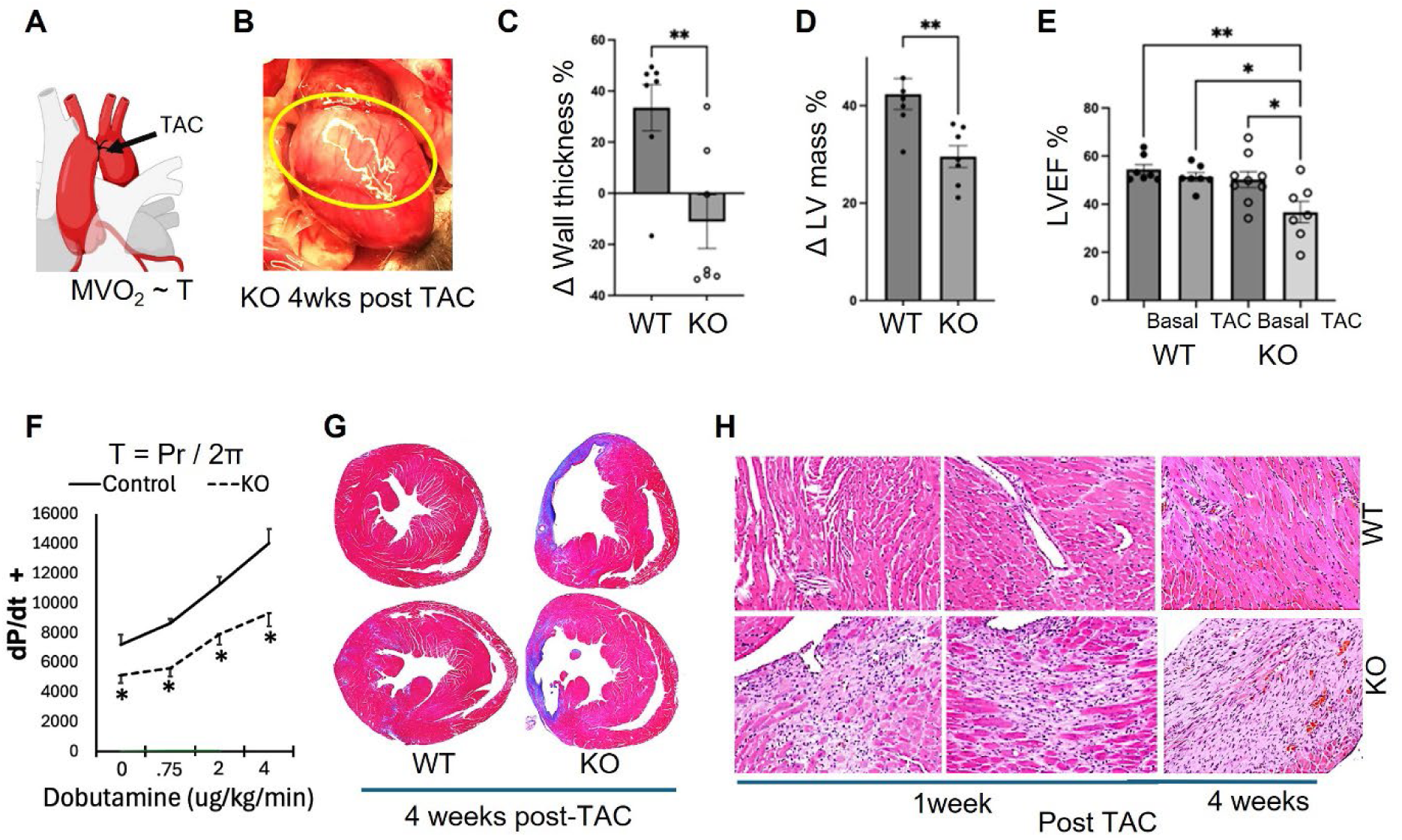
Renalase is required for adaptive cardiac hypertrophy. A. Transverse aorta constriction (TAC) induces pressure (P) overload on the left ventricle (LV), thus increasing myocardial wall tension (T) and myocardial oxygen consumption (MV02) per the LaPlace relationship. Compensation for increased T and consequent dilation of the LV (r = increased radius of curvature) involves thickening of the LV wall (hypertrophy = π) – an essential and fundamental biological response involving protein synthesis and increased myofibrils in cardiomyocytes. B. Renalase mice had a defective hypertrophic response to TAC, with 50% higher mortality, loss of myocytes, and aneurysmal bulging of the LV free wall (yellow oval). C-E. Echocardiography before and 4 weeks after TAC demonstrated increased LV wall thickness in control wild type (WT) mice, but thinning in RNLS KO hearts, an attenuation of the compensatory increase in LV mass, and a decreased ejection fraction (LVEF) in RNLS KO mice (E). F. Hemodynamic assessment with a transducer-tipped microcatheter during infusion of increasing doses of the inotropic agent dobutamine four weeks after TAC. The reduced LVEF and the reduced rate of rise of pressure during contraction (dPdt) of the KO hearts indicate reduced contractility in KO hearts post-TAC. *p<0.05. G. Trichrome staining 4 weeks post-TAC demonstrates severe thinning of the LV free wall, along with scar formation in the KO hearts H, Histological analysis of WT and KO hearts and 1 and 4 weeks after TAC. There was progressive cardiomyocyte loss, inflammation, thinning, and replacement scar tissue in the KO hearts from 1-4 weeks. Notably, TAC does not involve coronary arteries, and the areas of thinning did not correlate with a specific coronary territory.

## DISCUSSION

The secretory protein renalase (RNLS) has broad protective effects in transformed normal and cancer cells in culture and in vivo models of injury and cancer ^8,9,47–50^. For example, during cisplatin or ischemia-induced AKI, extracellular RNLS is crucial for the recovery of kidney function after the initial insult ^8^.

Intracellular RNLS is an NADH oxidase ^18,19^. We have reported that RNLS deletion results in a steady-state fall of > 50% in cellular ATP ^7^. In a recent study, we used RNAseq to characterize differentially expressed mRNAs in injured kidneys of wild-type or RNLS-deficient mice. RNLS (or RNLS treatment) upregulated mitochondrial genes, suggesting enhanced mitochondrial biogenesis ^8^.

Our study confirms and expands on these findings, providing deep insights into the intracellular mechanisms by which RNLS exerts its protective effects. We discovered that RNLS localizes to mitochondria and nuclei, suggesting roles beyond extracellular signaling. Our central finding is that RNLS can bind to mitochondrial ATP synthase, reprogram mitochondrial and cellular metabolism, and orchestrate a specific translational and subsequent transcriptional program that promotes cell survival.

A pivotal observation of our study is that extracellular RNLS induces a sustained increase in protein synthesis rate in serum-deprived cells. Notably, this increase occurs without a corresponding increase in cell proliferation, indicating a selective enhancement of cellular protein content. Adding extracellular RNLS increases intracellular RNLS levels for up to 24 hours. This positive feedback loop may implicate extracellular RNLS signaling through STAT3, a regulator of RNLS transcription ^51,52^.

Our proteomic analysis revealed that RNLS modulates cellular metabolism (oxidative phosphorylation, TCA cycle, glycolysis), the most enriched biological processes at one hour. These changes are accompanied by increased ribosomal biogenesis and translation of specific proteins, as evidenced by upregulation of several components of the 40S subunit (Rps7, Rps8, Rps10, Rps11, and Rps18). The pentose phosphate pathway and transcription become most enriched later. These data suggest that RNLS induces cellular metabolism changes that promote selective mRNA translation followed by gene transcription. An immunoblot of the puromycin cell lysate after RNLS treatment suggested a marked increase in puromycin uptake by the cells, indicative of an increased protein synthesis rate. The protein synthesis rate was decreased by adding Dex or cyclosporin A to cells before the puromycin addition, confirming the direct regulation of protein synthesis rate by ATP synthase leak modulation ^23^, and this was further confirmed by the absence of an increase in protein synthesis rate upon addition of exogenous RNLS to RNLS KO cells. Blocking RNLS signaling with Caloxin 1b, a selective inhibitor of the RNLS receptor PMCA4b, decreased protein synthesis rate in RNLS-expressing WT cells, suggesting that receptor activation increases intracellular RNLS activity. PKA-selective inhibitors also decreased the RNLS-enhanced protein synthesis rate. We hypothesize this could be partly due to dephosphorylation and activation of inhibitory factor 1 (IF1). IF1 prevents ATP hydrolysis ^53^, the latter a required element of leak metabolism ^23,54^. The p38 MAPK inhibitor also reversed the enhancement of the rate of protein synthesis by RNLS, suggesting an alternative mechanism through which RNLS increases the protein synthesis rate in RNLS-expressing cells. These findings suggest a sophisticated mechanism through which RNLS selectively regulates protein synthesis to optimize cellular stress response.

RNLS is highly expressed in renal proximal tubular cells. Advanced microscopy techniques showed that RNLS is concentrated in the mitochondrial matrix of various cell types, including renal proximal tubular cells. This unexpected localization suggested a mitochondrial function for RNLS. Although it is a secretory protein, our studies indicate that RNLS selectively targets intracellular organelles such as mitochondria. RNLS has a signal sequence at its N-terminus and a predicted mitochondrial targeting signal at the C-terminal region of the signal sequence. Regulated bimodal targeting of proteins with ER and mitochondrial targeting sequences has been reported for cytochrome p450 and could explain mitochondrial localization in some cell types. The details of RNLS’s cellular pathway, including whether it undergoes regulated targeting to the ER or mitochondria, are currently under investigation.

The localization of RNLS to mitochondria and the dramatic reduction in cellular ATP seen after its genetic deletion suggest that it may directly impact steps in energy production. The question remains whether the ATP synthase is partially running in reverse to help maintain the mitochondrial membrane potential to support leak metabolism. Our studies show that mitochondrial complex I and complex II activities decreased in isolated mitochondria from the RNLS KO. Treatment of isolated WT or RNLS KO mitochondria with full-length RNLS, but not with a peptide containing the RP220 site, increased Complex I and Complex II activities, suggesting a direct role for RNLS in supporting the rate of electron transport and maintaining the mitochondrial membrane potential.

A crucial insight from our study is the discovery that RNLS interacts with the mitochondrial ATP synthase F1 subunits, triggering the opening of the ATP synthase c-subunit leak channel. Although the effects of RNLS on Complex I could be due to its NADH oxidase activities, it is unlikely that this could account for its enhancement of Complex II activity. Therefore, we investigated RNLS’s effects on the ATP synthase. Proximity ligation analysis showed that mitochondrial RNLS and the ATP synthase F1 were separated by 40 nm or less, a value that would allow molecular interaction. Further studies showed that RNLS was near the ATP synthase F1 alpha and beta subunits, not the stator protein, OSCP. This supports a role for RNLS in regulating ATP synthase leak channel activity.

ATP synthase is another determinant of mitochondrial membrane potential, in addition to electron transport. Uncoupling of oxidation from phosphorylation by the opening of an inner mitochondrial membrane leak channel within the ATP synthase membrane embedded portion is directly causative of “leak metabolism” whereby the ATPase c-subunit leak channel opening drives an aerobic glycolytic metabolism associated with high glucose uptake, high lactate production, elevated TCA cycle enzymes and reversal of the ATP synthase enzyme to hydrolyze, rather than synthesize, ATP ^23,39,55–57^. The ATP synthase c-subunit ring forms a voltage-dependent uncoupling channel, the inactivation of which occurs through the binding of the F1 subcomplex to the c-subunit ^33,34,58,59^. Physiologically, this metabolism is required for early embryonic development and cell fate determination ^60^. High c-subunit levels are also found at later developmental periods, *e.g.,* just before the period of rapid synaptogenesis in the brain, and persistently dysregulated c-subunit expression after this critical period contributes to autism-associated behaviors in the Fragile X Syndrome animal model and to aberrantly elevated protein synthesis rates in murine neurons and patient cells^23^.

Endogenous and pharmacological regulators bind to the F1 subcomplex, altering the conformation of the complex to activate or inactivate the channel. ^33,61^. In the current study, we found that RNLS interacted with the F1 to cause a rapid and profound increase in the ATP synthase leak channel activity. The addition of specific and selective pharmacological inhibitory reagents to the patch recording showed that the RNLS-induced channel activity was likely to be caused by opening of both the ATP synthase leak channel (inhibited by Dex^38^) and the opening of the free c-subunit leak channel (inhibited by ATP) ^34^. We found that the aerobic glycolytic leak metabolism caused by c-subunit leak channel opening produced the increased protein synthesis rate that plausibly aided in kidney tubule cell repair and repair of damaged mitochondria, consistent with our previous report that showed an increase in transcripts indicative of mitochondrial biogenesis ^8^. Glycolytic proteins were upregulated in our proteomics search. To confirm aerobic glycolysis, we analyzed oxygen consumption in primary kidney cells isolated from WT and RNLS KO mice and demonstrated increased oxygen uptake in the presence of RNLS, consistent with activation of an aerobic state. Immunoblots of HKII, GPI, and TKT are also supportive of this glycolytic metabolism.

Electron micrographs obtained of the kidney proximal tubule S3 region at 4 days after cisplatin AKI confirmed that RNLS was required for normal proximal renal epithelial cell repair. In the WT injured kidney, we documented a high percentage of immediately adjacent membrane ER and mitochondria membranes, and we also noted abundant lipid-containing cytoplasmic vesicles, consistent with active synthetic and repair processes ^62^. We also found that cisplatin injury resulted in an increased number of small mitochondria, consistent with either mitochondrial fission during repair and/or mitochondrial biogenesis. PGC1alpha protein level was consistent with active mitochondrial biogenesis ^63^. In the RNLS KO cells, in contrast, autophagy and mitophagy markers were increased, whereas PGC1alpha was decreased, suggesting aberrant mitophagy instead of repair ^64^. The electron micrographs of the RNLS KO injured kidney showed a reduction in the close approximation of ER with mitochondria membranes and a striking decrease in mitochondrial numbers, consistent with increased Parkin-dependent mitophagy ^65^.

Cardiac hypertrophy is a fundamental and essential biological response to physiological and pathological stimuli ^66^. When faced with an increased workload, such as during exercise, pregnancy, or hypertension, the terminally differentiated cardiac myocytes of the adult heart undergo hypertrophy through increased protein synthesis and myofibril formation. RNLS likely contributes to these adaptive changes. Notably, in TKPTS cells exposed to RNLS for 24 hours, tropomyosin 1 (TPM1) protein expression increased by ∼19-fold (Supplemental Table 1). TPM1 is critically important for cardiac muscle contraction and adaptive hypertrophy, and disease-causing variants are linked to severe cardiac hypertrophy^67^. We show that RNLS KO mice have a defective cardiac hypertrophic response to chronic pressure overload, characterized by fibrosis, thinning, aneurysmal bulging of the left ventricular free wall, and a ∼50% rate of cardiac rupture in 4 weeks. Compared to sedentary healthy volunteers, those who completed an eight-week aerobic or anaerobic exercise or strength training program showed a significant increase in their plasma RNLS ^68^. Low plasma RNLS portends worse cardiovascular outcomes and increased mortality for patients with acute organ injury ^69,70^. In animal models of acute organ injury, administration of exogenous RNLS prevents severe injury and improves mortality ^7,8,11,47^

In summary, RNLS is localized to mitochondria, where two functions have been identified. First, it stimulates the activities of Complex I and Complex II, affecting electron transfer and cellular energetics. Second, it binds to mitochondrial ATP synthase, opening its leak channel. This induces leak metabolism and an increased protein synthesis rate to repair damaged cells and organelles, including mitochondria. This repair process reduces mitophagy and inhibits the death of the proximal tubule cells. Although the mechanism by which mitochondrial leak enhances protein synthesis rate is as yet unknown, it is likely to occur through changes in the phosphorylation state of RNA-binding proteins in non-membrane-delimited RNA-containing granules ^71,72^ close to mitochondria. The electron micrographic images of proximal tubule cells demonstrate the proximity of the rough and smooth ER to mitochondria, supporting this notion. In conclusion, we have identified and characterized an RNLS-dependent molecular mechanism of mitochondrial and protein synthesis-dependent response to stressful stimuli that promotes mitochondrial health and physiological adaptation and induces injury repair.

## MATERIALS AND METHODS

### STAR★Methods

Key resources table.

Key resources table
| REAGENT or RESOURCE | SOURCE | IDENTIFIER |
| --- | --- | --- |
| **Antibodies** |  |  |
| Mouse anti-Puromycin | Millipore | MABE343MI |
| Mouse anti-Puromycin [3RH11] | Kerafast | Equation 0001; RRID:AB_2620162 |
| Rabbit anti-b-Actin | Cell Signaling | 8457 |
| Rabbit anti-RNLS, monoclonal [m28] | in house production | N.A. |
| Goat anti-RNLS | abcam | ab31291 |
| GPI | Proteintech | 15171-1-AP |
| HK-II | Cell Signaling | 2867 |
| TKT | Cell Signaling | 64414 |
| Rabbit anti-COX IV | Cell Signaling | 4850 |
| Tom20 | Cell Signaling | 42406 |
| podocin | Sigma-Aldrich | P0372 |
| Mouse anti-ATP5A | abcam | ab14748 |
| Mouse anti-ATP5B | Santa Cruz Biotechnology | sc-55597 |
| Mouse anti-ATP5O | Santa Cruz Biotechnology | sc-365162 |
| Rabbit anti-PGC1a | Novus Biologicals | NBP1-04676 |
| Rabbit anti-OPA1 | Cell Signaling | 80471 |
| Rabbit anti-p-Thr172-AMPKa | Cell Signaling | 2535 |
| Rabbit anti-HO-1 | Cell Signaling | 43966 |
| Rabbit anti-Parkin | Cell Signaling | 32833 |
| Rabbit anti-LC3A/B | Cell Signaling | 12741 |
| Rabbit anti-DRP1 | Santa Cruz Biotechnology | sc-32898 |
| Mouse IgG | Cell Signaling | 87582 |
| Rabbit IgG | Cell Signaling | 49077 |
| HRP-linked anti-mouse IgG | Cell Signaling | 7076; RRID:AB_2099233 |
| HRP-linked anti-rabbit IgG | Cell Signaling | 7074; RRID:AB_2099233 |
| ATTO 647N linked anti-goat IgG | Rockland | 605-456-013 |
| ATTO 594 linked anti-rabbit IgG | Sigma-Aldrich | 77671 |
| AF594 linked anti-rabbit IgG | Cell Signaling | 8889 |
| AF488 linked anti-mouse IgG | Cell Signaling | 4408 |
| AF488 linked anti-rabbit IgG | Cell Signaling | 4412 |
| **Bacterial Strains** |  |  |
| One Shot TOP10 chemically competent Escherichia coli | Invitrogen | C404010 |
| BL21 (DE3) competent cells | Invitrogen | C600003 |
| **Chemicals** |  |  |
| Puromycin Dihydrochloride from Streptomyces alboniger | Sigma-Aldrich | P8833 |
| Bradford Protein Assay kit | BioRad | 500-0201 |
| Trypan Blue Solution, 0.4% | ThermoFisher Scientific | 15250061 |
| MitoTracker® Red CMXRos | Cell Signaling | 9082 |
| TMRM, tetramethylrhodamine, methyl ester | Invitrogen | T668 |
| Dex, Dexpramipexole Dihydrochloride | Sigma-Aldrich | SML0392 |
| Oligomycin, Seahorse XF Mito Stress Test Kits | Agilent | 103016-100 |
| FCCP, Seahorse XF Mito Stress Test Kits | Agilent | 103016-100 |
| AA, Antimycin A, Seahorse XF Mito Stress Test Kits | Agilent | 103016-100 |
| Rot, Rotenone, Seahorse XF Mito Stress Test Kits | Agilent | 103016-100 |
| Rapamycin | Cell Signaling | 9904 |
| Cyclosporine A | Caymen Chemical | 12088 |
| Caloxin 1b1 | AnaSpec | AS-64236 |
| H89 | Selleckchem | S1582 |
| SB203580 | Cell Signaling | 5633 |
| 666-15 | Selleckchem | S8846 |
| Lipofectamine 3000 transfection reagent | Thermo Fisher | L3000001 |
| cOmplete, Mini, EDTA-free Protease Inhibitor Cocktail | Roche | 11836170001 |
| phosphatase inhibitors | Sigma-Aldrich | P5726 |
| Protein A magnetic beads | Cell Signaling | 73778 |
| RIPA buffer (10x) | Sigma-Aldrich | R0278 |
| formaldehyde | Sigma-Aldrich | 47608 |
| acrylamide | Sigma-Aldrich | A9099 |
| VectaShield with DAPI | Vector Laboratories | H-1200 |
| **Critical commercial assays** |  |  |
| pan-expansion microscopy (PanExM) | Panluminate Inc | https://www.panluminate.com/ |
| Seahorse XFe96 FluxPak | Agilent Technologies | 102416-100 |
| Duolink® Proximity Ligation Assay | Millipore-Sigma | DUO94004 |
| Mitochondria isolation kit | Qiagen | 37612 |
| NADP/NADPH-Glo™ Assay kit | Prmega | G9081 |
| Stanbio urea nitrogen direct kit | Fisher Scientific | SB-0580-250 |
| **Deposited data** |  |  |
| See Excel file named "Datasets of mass spec" in the manuscript prep folder. |  |  |
| **Experimental models: Cell lines** |  |  |
| TKPTS | ATCC | CRL-3361; RRID:CVCL_UJ13 |
| HEK293T | ATCC | CRL-11268; RRID:CVCL_1926 |
| Hela | ATCC | CRM-CCL-2; RRID:CVCL_0030 |
| MiaPaCa2 | ATCC | CRM-CRL-1420; RRID:CVCL_0428 |
| **Experimental models: Mouse** |  |  |
| C57Bl/6j as wild type | Jackson Laboratory | 000664 |
| RNLS KO | in house | N.A. |
| **Recombinant DNA** | This study | N.A. |
| pET28-RNLS | This study | N.A. |
| pcDNA3.1-RNLS | This study | N.A. |
| 1-66 GFP (peGFP3n-1-66RNLS) |  |  |
| **Peptides and Recombinant Proteins** | LifeTein | customer synthesized |
| RP10 | LifeTein | customer synthesized |
| scRP | This study | N.A. |
| RNLS | This study | N.A. |
| **Software and algorithms** |  |  |
| Prism 10 | GraphPad (commercial) | https://www.graphpad.com |
| imageJ version 1.54p | U. S. National Institutes of Health | https://imagej.net/software/fiji/downloads |
| Scaffold 5.3.4 | Proteome Software Inc. | https://www.proteomesoftware.com/ |
| Photoshop 2024 | Adobe Inc. | https://adobe.com |
| Qluore Omics Explorer | Qluore, Lund, Sweeden | https://qluore.com/ |
| STRING | STRING Consortium 2025 | https://string-db.org/ |

Resource availability. Further information and requests for resources and reagents should be directed to and will be fulfilled by the lead contact, Dr. Desir.

#### Materials availability

Plasmids, cell lines, and animal lines generated in this study will be available upon request. However, if there is potential for commercial application, we may require payment and/or a completed Materials Transfer Agreement.

#### Data and code availability

The lead contact will share all data reported in this paper upon request. Original images are available at Links, or accession numbers are also listed in the key resources table. The datasets are publicly available. This paper does not report the original code. Any additional information required to reanalyze the data reported in this work paper is available from the lead contact upon request.

### Experimental model and study participant details

#### Animal maintenance and AKI mouse model

Mice were maintained at Veterans Affairs Medical Center (VAMC), West Haven, CT. All experimental procedures were conducted according to the guidelines and regulations for animal care and use by the Institutional Animal Care and Use Committee of the VAMC, and the authors complied with the ARRIVE guidelines. Mice were kept under a 12-h day/night cycle with food and water provided ad libitum. All experiments were repeated on at least two separate occasions. We used the ARRIVE reporting guidelines to report the animal studies.

Homozygous RNLS KO mice were generated as previously described. ^11^. The KO mice have been generated by backcrossing with WT mice (Strain C57BL6J, Jackson Laboratory, Cat# 000664) for ten generations to preserve the C57BL6J background. Both genotyping and mRNA levels confirmed the disruption of the RNLS gene in the KO mice. To establish AKI, adult WT, and RNLS KO mice were administered a single dose of CP (15 mg/kg) subcutaneously. Control mice were administered saline. Four days post-CP, the animals were sacrificed, and blood was collected in a microtube precoated with heparin (H3393, Sigma) and centrifuged at 2,000 g for 10 min. The upper phase after centrifuging was collected as plasma. Kidneys were collected for further analysis.

#### Cell culture and cell counting

Immortalized mouse proximal tubule cell line TKPTS (CRL-3361, RRID: CVCL_UJ13), HEK293T cells (CRL-11268; RRID: CVCL_1926), Hela cells (CRM-CCL-2, RRID: CVCL_0030), and MIA PaCa-2 (CRM-CRL-1420, ATCC) were obtained from ATCC. Mycoplasma negative status was confirmed by PCR Detection Kit. These cells were cultured in a humidified incubator at 37 °C with 95% air and 5% CO2 on culture dishes (Corning). The culture medium used for each cell line was followed by ATCC instruction. Upon reaching 90% of confluency, cells were PBS-washed, trypsinized, and then split at 1:5 (TKPTS and MiaPaCa2 cells) to 1:10 (293T and HeLa cells). Cells were frozen in liquid nitrogen for long-term storage with freezing media (FBS + 10% DMSO). Cells were used for up to 10 passages before a new aliquot was thawed.

TKPTS cells were cultured in serum-free medium for 24 hours and then incubated with either scRP, inactive RNLS agonist, or with rRNLS. The cells were harvested 24 hours later, stained with Trypan Blue (15250061, Thermo Fisher Scientific) staining, and live cells were identified and counted by microscopy in the hemocytometer (22-600-114, Fisher Scientific).

Primary culture of renal cells was isolated from kidneys of WT and RNLS KO animals according to previously described method^73^.

#### Recombinant RNLS protein and synthetic RNLS peptide

Recombinant RNLS proteins were prepared as described ^74^. Briefly, DNA encoding human RNLS-1 was synthesized and cloned into a pET28a vector with an HIS tag at the N-terminal of RNLS. The construct was transformed into bacterial strain BL21 (DE3) to over-express RNLS. RNLS was purified using HisTALON™ Gravity Column Purification Kit (635654, Takara). RNLS peptide RP10 and its scrambled peptide (scRP) were custom synthesized by LifeTein: GKIDVPWAGQYITSNPAIRFVSIDNKKRNIESSEIGP and scRP sequence, respectively.

#### RNLS constructs and DNA plasmid transfections

Human RNLS in pET28a is subcloned into pcDNA3.1 for mammalian expression. N-terminus of RNLS (nt1-66) fused to GFP (pEGFP-N3) was constructed. According to the manufacturer’s specifications, HeLa or HEK293T cells were transfected pcDNA3.1c-RNLS (OriGene) using Lipofectamine 3000 reagent (L3000015, Thermo Fisher Scientific).

#### Drug Treatments

To test the effect of RNLS in protein synthesis, TKPTS cells were serum-starved for 24 h, followed by incubation with recombinant RNLS or RP10 or scRP at 50 µg/ml for the indicated time. Puromycin was added to the cells at 10 µg/ml 10 minutes before cell lysis. In some experiments, a serial concentration (0 – 100µM) of Dexpramipexole Dihydrochloride (Dex, SML0392, Sigma-Aldrich), 0-500nM Cyclosporin A (CsA, Item No. 12088, Cayman Chemical), 0-100µM ATP2b4 inhibitor caloxin1b1 (Caloxin, AS-64236, AnaSpec), 0-500nM mTOR inhibitor rapamycin (Cat# S1039, Selleck Chemicals), 1µM H89 (S1582, Selleckchem), which is a selective PKA inhibitor that binds to the catalytic subunit of protein kinase A and inhibits its action, 10 µM SB203580 (#5633, Cell Signaling Technologies), which inhibits p38 catalytic activity by binding to the ATP binding pocket and also inhibits the phosphorylation and activation of protein kinase B (PKB, also known as Akt), or 20µM CREB inhibitor 666-15 (S8846, Selleckchem) was added to the cells 30 min before the addition of RNLS or RNLS peptides.

#### Western Blotting

Cells were lysed with ice-cold RIPA buffer [RIPA (R0278, Sigma Aldrich) supplemented with protease inhibitor cocktails (11836170001, Roche) and phosphatase inhibitors (P5726, Sigma). Kidney tissue collected from mice was left in RIPA buffer and immediately homogenized in a BeadBug homogenizer (RS7020, Benchmark Scientific) at 4000 rpm for 100 sec. After centrifuging to remove debris, the protein concentration in the resulting supernatant was determined using the Bradford Protein Assay (500-0201, BioRad). The supernatant was mixed with the Reducing Laemmli Sample Buffer (1610747, BioRad) and heated at 95 °C for 5 minutes. Subsequently, 10-20 µg of proteins were loaded onto a Criterion TGX precast gel (5671033, BioRad) and separated with SDS running buffer (1610744, BioRad). The separated proteins were later transferred onto a PVDF membrane (1620177, BioRad) and blocked with EveryBlot Blocking Buffer (12010020, BioRad). The membranes were then incubated with primary antibodies diluted in the Blocking Buffer overnight, followed by secondary antibodies dissolved in the Blocking Buffer. Proteins were detected using a chemiluminescent substrate (34580, Thermo Fisher Scientific). Signals were stripped in Restore™ Western Blot Stripping Buffer (21059, Thermo Fisher Scientific) for re-blotting. ImageJ software quantified protein bands (U. S. National Institutes of Health, Bethesda, MD, USA). Each band’s intensity was normalized to the intensity of the housekeeping protein β-Actin band.

#### Immunoprecipitation of puromycin-labelled peptides

A 100 µg of protein lysate from cell cultures treated with reagents and puromycin was incubated at 4°C overnight with 1μg of anti-puromycin antibody (Kerafast). Next, 20 µL of protein A magnetic beads (73778, Cell Signaling) were added to the samples for overnight incubation at 4°C. Beads were washed 3 times with 1x RIPA buffer and processed for mass spectrometry analysis.

#### Mass Spec

Following immunoprecipitation, pull-down proteins were eluted in Laemmli Sample Buffer, boiled, and purified in a stacking gel of SDS-PAGE; proteins in the gel were excised for bottom-up protein identification by LC/MS/MS. Gel bands were prepared as described. ^75^ Briefly, excised gel bands in 1.5 Eppendorf tubes are washed 4 times; first with 500 μL 60% acetonitrile containing 0.1%TFA and then with 5% acetic acid, then with 250 μL 50% H2O/50% acetonitrile followed by a 250 μL 50% CH3CN/ 50 mM NH4HCO3, and a final wash with 250 μL 50% CH3CN/10 mM NH4HCO3 before removal of wash and complete drying of gel pieces in a Speed Vac. 10 μL of a 0.1 mg/mL stock solution of trypsin (Promega Trypsin Gold MS grade) in 5mM acetic acid is freshly diluted into a 140 μL solution of 10mM NH4HCO3 to make the working digestion solution. 124 μL of the working digestion solution is added to the dried gel pieces (an additional 10 mM NH4HCO3 was added to ensure gel pieces are completely submerged in the digestion solution) and incubated at 37 °C overnight. The sample is then stored at −20 °C until analysis. Tryptic peptides were separated on a nanoAcquity™ UPLC™ column (Waters) coupled to a Q-Exactive Plus mass spectrometer. High-resolution tandem LC-MS/MS data were collected by Higher-Energy Collisional Dissociation (HCD) with a 1.4 Da window, followed by a normalized collision energy of 32%. Resulting LC-MS/MS data were analyzed and processed through Proteome Discoverer (v.2.2, linked to MASCOT search engine v.2.4) and further integrated with Scaffold (v.4.8, Proteome Software Inc.).

#### Pan-expansion microscopy (pan-ExM) of mouse kidneys

Kidney tissue experiments were conducted in adult male mice in WT (C57BL/6j). The mice were anesthetized with isoflurane, transcardially perfused first with ice-cold 1× PBS and then with 4% formaldehyde plus 20% acrylamide (A9099, Sigma) in 1× PBS, pH 7.4. Kidneys were isolated and post-fixed 4-6 hours in the same perfusion solution at 4 °C. Kidneys were subsequently washed and stored in PBS at 4 °C. Kidneys were mounted in ice-cold PBS and sectioned at 70 µm using a vibrating microtome (Vibratome 1500, Harvard Apparatus). Kidney slices were selected and washed four times for 15 min in PBS at room temperature. Sections were stored in PBS at 4 °C for up to 3 months. Kidney sections were subjected to pan-ExM as previously described. ^76^. Kidney gels processed with pan-ExM were incubated overnight with 1:300 diluted goat-anti-RNLS antibody (ab31291, Abcam) or 1:250 diluted rabbit-anti-podocin (P0372, Sigma) in antibody dilution buffer (0.05% TX-100 + 0.05% NP-40 + 0.2% BSA in 1× PBS) on a rocking platform at 4 °C. Next, samples were incubated with donkey anti-goat at 1:375 (605-456-013, Rockland) or ATTO 594 labeled goat anti-rabbit (77671, Sigma), respectively. Gels were then washed in 0.1% (v/v) TX-100 in 1× PBS four times for 30 min to 1 h each on a rocking platform at RT and once overnight at 4 °C. The gels were subsequently washed in 0.1% (v/v) TX-100 in 1× PBS four times for 30 min to 1 h each on a rocking platform at RT and once overnight at 4 °C. The gels were mounted on glass-bottom dishes (35 mm; no. 1.5; MatTek). A clean 18-millimeter diameter coverslip (0117580, Marienfeld) was put on top of the gels after draining excess water using Kimwipes. The samples were then sealed with two-component silicone glue (Picodent Twinsil, Picodent, Wipperfürth, Germany). After the silicone mold hardened (typically 15–20 min), the samples were stored in the dark at 4 °C until imaged. All confocal images were acquired using a Leica SP8 STED 3X with a SuperK Extreme EXW-12 (NKT Photonics) pulsed white light laser as an excitation source. Images were acquired using either an HC FLUOTAR L 25×/0.95 NA water objective, APO 63×/1.2 NA water objective, or an HC PL APO 86×/1.2 NA water CS2 objective. Application Suite X software (LAS X; Leica Microsystems) was used to control imaging parameters. ATTO594 was imaged with 585-nm excitation. Images were visualized, smoothed, gamma-adjusted, and contrast-adjusted using FIJI/ImageJ software. STED and confocal images were smoothed for display with a 0.4 to 1.5-pixel sigma Gaussian blur. Minimum and maximum brightness were adjusted linearly for optimal contrast. All line profiles were extracted from the images using the Plot Profile tool in FIJI/ImageJ.

#### Immunofluorescence staining

MiaPaCa cells were plated on coverslips (174969, Thermo Fisher Scientific) and fixed in 4% paraformaldehyde for 15 min at RT and permeabilized in buffer containing 0.05% saponin. Cells were incubated with rabbit anti-RNLS, clone m28 antibody, and CoxIV antibody (4850, Cell Signaling Technologies) overnight at 4 °C, followed by incubation with AF594-linked anti-rabbit IgG and AF488-linked anti-mouse IgG. The cover slip was mounted with VectaShield with DAPI, and cells were imaged on a Nikon A1R multiphoton confocal microscope with NIS Elements software. In some experiments, m28 was preincubated with RNLS peptide RP10 (5:1 peptide to antibody) before incubation with cells to show m28 specificity.

HEK293 cells transfected with the N-terminus of RNLS (nt1-66) fused to GFP (pEGFP-N3) were fixed in 4% paraformaldehyde for 15 min at RT and permeabilized in buffer containing 0.1% Triton X-100. Cells were incubated with Tom20 antibody (42406, Cell Signaling) for 2h at RT, followed by incubation with AF594-linked anti-rabbit IgG. The cover slip is mounted with VectaShield with DAPI, and cells were imaged on a Zeiss Axiophot microscope.

HeLa cells stably transfected with pcDNA3.1-RNLS and renal cells isolated from WT or RNLS KO mouse kidneys were grown on coverslips to reach the confluence of 70% followed by incubation with 200nM MitoTracker® Red CMXRos (9082, Cell Signaling) for 30 min at 37°C. Cells were fixed in ice-cold methanol for 15 minutes at –20°C. Cells were incubated with 0.35ug/ml m28 overnight at 4^0^C, followed by AF488-linked anti-rabbit IgG (A-11008, Thermo Fisher Scientific). The coverslips were mounted with VectaShield with DAPI (H-1200, Vector Laboratories). Cells were imaged on a Leica SP8 confocal microscope with a 63X/1.2 NA W objective.

#### Measuring activities of Complexes I and II

Mitochondria were isolated and purified from the kidneys of WT and RNLS KO mice using a Qproteome™ Mitochondria isolation kit (37612, Qiagen), according to the manufacturer’s protocol. Mitochondria were incubated with 200nM tetramethylrhodamine ethyl ester (TMRE) in the presence of Complex I substrates 4mM pyruvate and 4mM malate, or Complex II substrate 4mM succinate. The fluorescence intensity of TMRE was measured in a plate reader (Model FLx800, BioTek Instruments). In some experiments, rRNLS was included in the reaction at 50µg/ml.

#### Proximity Ligation Assay (PLA)

HeLa cells were seeded on poly-lysine-treated slides and transfected with pcDNA3.1c-RNLS on the following day. 36 hours after the transfection, cells were permeabilized for 15 minutes with 0.5% Triton X-100 in 1X PBS and blocked with 4% goat serum for 1 hour at 37°C. Then, PLA was performed following the procedure described in a previous publication (Dieck et al., 2015; Sambandan et al., 2017) using PLA PLUS Probe Anti-Rabbit RNLS and PLA MINUS probe anti-mouse ATP5A (or anti-mouse ATP5B, anti-mouse ATP5O). After the ligation and the amplification process, a Far Red Duolink® PLA Fluorescence Detection Reagent kit was used to detect the interacting protein complex. Z-stack images were taken using a Zeiss Airyscan 880. The puncta, representing a direct interaction between the two interested proteins, were analyzed using Image J. An anti-Rabbit IgG and anti-Mouse IgG combination was used as the negative control.

#### Mitoplast Electrophysiology

Mitochondria were isolated from mouse liver or kidney using Qproteome Mitochondria mitochondrial isolation kit (Qiagen, Cat. No. 37612). In brief, cells were transferred to ice-cold isolation buffer, supplemented with 1x Halt protease inhibitor. Cells were minced, homogenized with a Dounce homogenizer, and centrifuged at 1000 × g to pellet nuclei, cell debris, and unbroken cells. The supernatant was centrifuged at high speed (6000 × g for 15 min at 4 °C); the pellet containing mitochondria was washed in isolation buffer and pelleted by centrifugation at 6000 × g. Protein concentration was determined by the BCA method using BSA as a standard. Isolated mitochondria were used immediately or stored at −80 °C until use. Mitoplasts were prepared by incubating 2 μl mitochondria in 30 mM KCl, 10 mM HEPES, pH 7.3, for 10 minutes, then centrifuging at high speed to wash and replacing the buffer with intracellular (recording) solution (120 mM KCl, 8 mM NaCl, 0.5 mM EGTA, 10 mM HEPES, pH 7.3). Each 2 μL was plated on glass-bottom dishes for visualization of mitoplasts that are resting and attached to the dish bottom. Recording electrodes were pulled from borosilicate glass capillaries (WPI) with a final resistance in the range of ∼50-100 MΩ. The patch-clamp recordings of kidney or liver mitoplasts were performed by forming a giga-ohm seal on the mitoplast membrane (mitoplast attached) using an Axopatch 200B amplifier (Axon Instruments) at room temperature (22–25 °C), then attempting to removing a patch from the mitoplast by quickly raising the patch electrode off the dish so that the matrix side of the mitoplast was bath-exposed. Recorded current signals performed in voltage clamp mode were filtered at 5 kHz using the amplifier circuitry. ATP (1 mM, final concentration), bongkrekic acid (BA, 5 μM), and Dexpramipexole (10 μM) were added into the bath during the recordings without perfusion. pCLAMP-10 software was used for electrophysiology data acquisition and analysis (Molecular Devices). All current measurements were adjusted for the holding voltage, assuming a linear current-voltage relationship: The resulting conductances are expressed in pS according to the equation G = I/V where G is conductance in pS, V is the membrane holding voltage in mV, and I is the peak membrane current in pA after subtraction of the baseline electrode leak current. Group data were quantified in terms of peak conductance during a 10-second sweep. All population data were expressed as mean ± SEM, where each N was a separate mitoplast recording.

#### Seahorse metabolic assay

According to the manufacturer’s protocol, OCR and ECAR were measured using a Seahorse Extracellular Flux Analyzer (XFe96). In brief, tubule cells were isolated from the kidneys of WT or RNLS KO mice as described^11^ and seeded in a Seahorse XFe96 Cell Culture Microplate (34222, Agilent) and grown to 60-70% confluency at the time of measurement. Agilent Seahorse Extracellular Flux Assay Kits were used for OCR and ECAR measurements. Initially, probes were hydrated in water in a non-CO2 37 °C incubator overnight, followed by a 1-hour incubation with a pre-warmed calibrant. The Seahorse XF DMEM medium (103575-100, Agilent) supplemented with 4.5g/L glucose, 1 mM sodium pyruvate, Glutamax, NEAA, and antibiotics (but excluding FBS) was substituted for regular culture media 1 hour before measurement in a 37 °C CO2-free incubator. Mitochondrial electron transport chain inhibitors were loaded into the probe cartridge. After probes were calibrated in the calibrant buffer, the cell culture plate was loaded for OCR and ECAR measurements.

#### Measurement of NADP and NADPH levels

TKPTS cells were plated in a 96-well plate, serum-starved for 24 hours, then treated with 1.35 µM rRNLS for the indicated time points. NADP and NADPH levels were measured separately in the same wells using the NADP/NADPH-Glo™ Assay kit (Promega, Cat # G9081) according to the manufacturer’s instructions. The protein concentration of the lysate was determined by Bradford assay using Protein Assay Dye (BioRad, cat # 500-0201). NADP and NADPH levels were normalized to total protein content.

#### Measurement of plasma creatinine and blood urea nitrogen (BUN)

Plasma creatinine levels were measured by mass spec at George M. O’Brien Kidney Center at Yale, and BUN levels were determined using a Stanbio urea nitrogen direct kit (Thermo Fisher Scientific, Waltham, MA, USA) according to the manufacturer’s instructions.

#### Kidney electron microscopy

Samples were fixed in 4% paraformaldehyde in 0.25M Hepes for 1 hour. Samples were rinsed in PBS, re-suspended in 10% gelatin, chilled, and trimmed into smaller blocks, and placed in a cryoprotectant of 2.3M sucrose overnight on a rotor at 4°C. They were transferred to aluminum pins and frozen rapidly in liquid nitrogen. The frozen block was trimmed on a Leica Cryo-EMUC6 UltraCut, and 65–75nm thick sections were collected using the Tokoyasu method. The frozen sections were collected on a drop of sucrose, thawed, placed on a nickel formvar/carbon-coated grid, and floated in a dish of PBS ready for immunolabeling. Grids were placed section side down on drops of 0.1M ammonium chloride to quench untreated aldehyde groups, then blocked for nonspecific binding on 1% fish skin gelatin in PBS. All grids were rinsed in PBS, fixed using 1% glutaraldehyde for 5 minutes, rinsed, transferred to a UA/methylcellulose drop, and then dried for viewing. Samples were viewed FEI Tencai Biotwin TEM at 80Kv. Images were taken using Morada CCD and iTEM (Olympus) software. Analysis of electron micrographs was performed using ImageJ software (NIH).

#### Cardiac pressure overload by aortic constriction

To induce pressure overload on the heart, we will use the murine TAC model, as previously described ^77^. Briefly, in anesthetized non-intubated mice, the transverse aorta is visualized via a small incision, a 7-0 nylon suture is tightened around the transverse aorta and a 27-gauge needle, and the needle is pulled free, resulting in a patent but significantly constricted aorta. Mortality is very low in WT mice, and TAC is optimized to balance survival and hemodynamic effects. Controls undergo sham surgery.

#### Statistical analysis

Statistical analysis was performed using Prism 10 (GraphPad Software, San Diego, CA). Data are presented as mean ± SEM. A paired or unpaired student’s two-tailed t-test was used to compare two groups. A one-way or two-way ANOVA test with Tukey post-hoc test was used for multiple comparisons. Statistical details and methods used in each experiment can be found in figures and figure legends. p<0.05 is considered statistically significant. p values are provided in figure legends (*p<0.05; **p<0.05; ***p<0.001; ****p<0.0001).

For mass spec result analysis, all experiments were performed in triplicate, and log2-transformed normalized protein abundance was analyzed and visualized using Qlucore Omics Explorer (v 3.10.9, Qlucore, Sweden). Proteins with more than 90% of missing values were removed. Differentially abundant proteins were determined by a two-sided unpaired t-test. Proteins were considered differentially abundant if p-values <0.05 and False Discovery Rate (FDR) q <0.05. Protein interaction network and clustering analysis were done on the STRING database (version 12.0, https://string-db.org/)^78^. Markov cluster algorithm with inflation parameter 3 was used to identify clusters of highly interconnected nodes and subsequent biological functions with FDR q <0.05.

## Acknowledgements

We thank the MS & Proteomics Resource at Yale University for providing the necessary mass spectrometers and the accompanying biotechnology tools funded in part by the Yale School of Medicine and by the Office of the Director, National Institutes of Health (S10OD02365101A1, S10OD019967, and S10OD018034). The funders had no role in study design, data collection and analysis, decision to publish, or preparation of the manuscript. We also would like to thank Jean Kanyo for assistance with mass spectrometry, sample preparation, and data collection.

Cellular and mitochondrial respiration (OCR) and acidification (ECAR) are assessed with the Agilent Seahorse XF Pro 96 by the Chemical Metabolism Core at Yale University. We thank Yale University’s Chemical Metabolism Core, especially Rebecca Cardone, for their assistance with the technique and contribution to this project.

George M. O’Brien Kidney Center at Yale, where the mouse plasma creatinine was measured, was funded by NIH grant P30 DK079310. We would also like to thank Lonnette Diggs for technical support.

We also thank Dr. Xinran Liu and Kimberly Zichichi (Yale Center for Cellular and Molecular Imaging) for assistance with electron microscopy and insightful scientific discussion.

This study was supported by Yale Research funds to GVD, NIH grants 045876 and 112706 to EAJ.

