## Supplemental Figures and Legends for "Renalase activates mitochondrial leak metabolism in response to cellular stress and to repair damage after injury"

### Table S1

Data sets of Mass spec, will be deposited to public domain.

### Figure S1

Pan-Expansion Microscopy of cortical (A) and glomerulus (B) of WT mouse kidney tissue. Protein density is displayed with a black-to-white scale. Arrows indicate structures of interest. (C) Podocin expression in the glomerulus. Pan-Expansion Microscopy of cortical WT mouse kidney tissue immunolabeled with anti-Podocin antibody. The positivity for Podocin is pseudocolored in a golden color in the merged image. Arrows indicate structures of interest. Scale bar = 1 μm.

### Figure S2

Specificity of m28 anti-RNLS monoclonal antibody and RNLS mitochondrial localization. (A) Immunostaining of RNLS using the m28 monoclonal antibody in renal cells isolated from WT or RNLS KO kidneys show positivity in WT renal cells only. (B) RNLS staining by m28 in MiaPaCa2 cells is prevented by preincubation with RNLS peptide. (C) RNLS contains a mitochondrial localization signal in its first 1-66 nucleotide. HEK293 cells transfected with the N-terminus of RNLS (NT1-66) fused to GFP (pEGFP-N3) show GFP in mitochondria. Scale bar = 2 μm.

### Figure S3

Quantification of mitoplast recordings, including WT (kidney or liver) and kidney RNLS KO (one-way ANOVA; *p=0.0381, *p=0.0359, **p=0.0019).

### Figure S4

(A) Principal component analysis of the 697 differentially abundant proteins at 1, 6, 12, and 24 hrs. of rRNLS treatment reveals minimal overlap between groups and unique protein expression profiles that characterize different time points. (B) KEGG enrichment pathway of RNLS effect on protein synthesis at the earliest measured time point of 1h. (C) Results from the STRING database clustering analysis (inflation parameter 3) showing the clusters and enriched pathways for each cluster at 1 hour (False discovery rate (FDR) $p<0.05$). (D-E) KEGG enrichment pathway for cluster 1 (D) and for cluster 2 (E).

**Figure S1**

**A. Proximal tubule mitochondria**

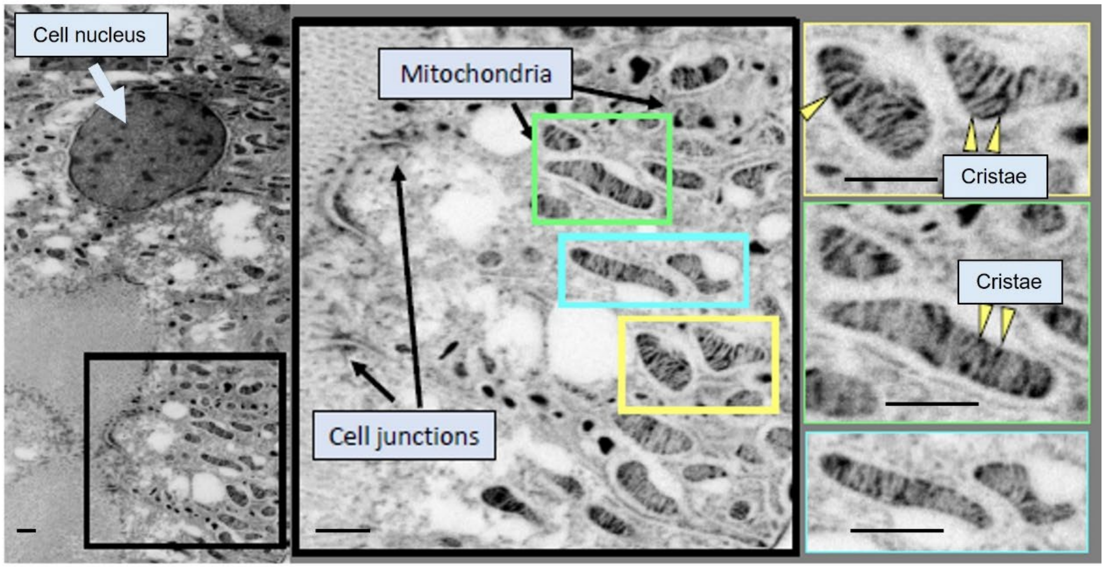


**B. Expansion microscopy of the glomerulus**

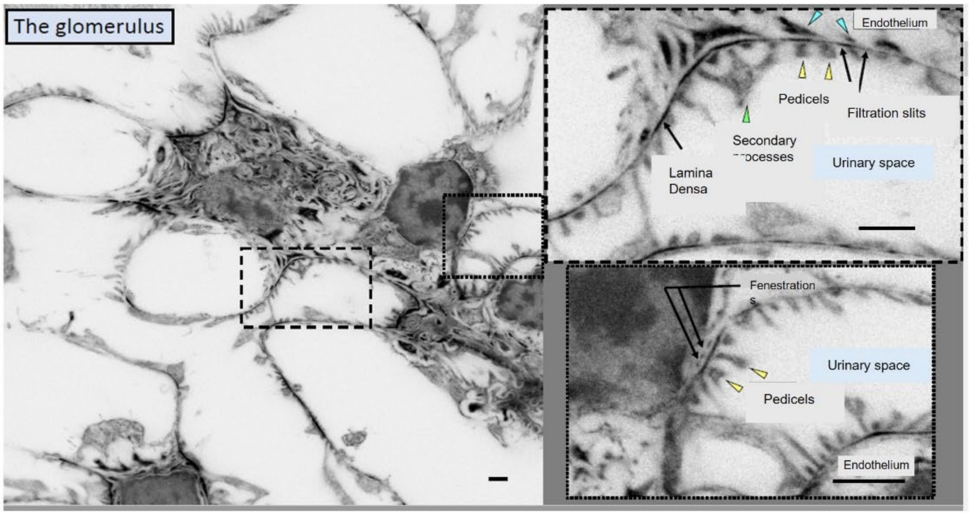


**C. Podocin expression in the glomerulus**

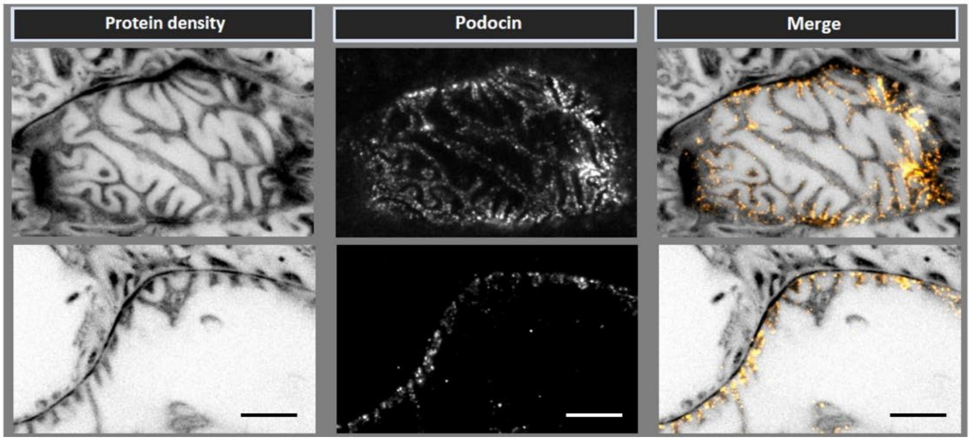

**Figure S2**

**A** **m28 is specific for RNLS**

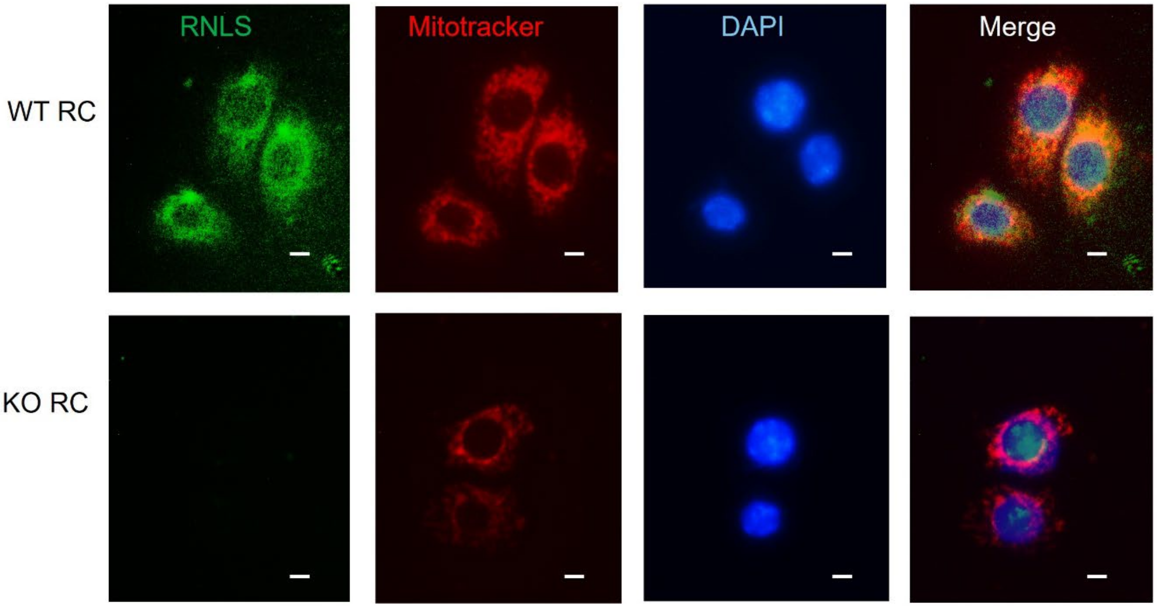


**B**

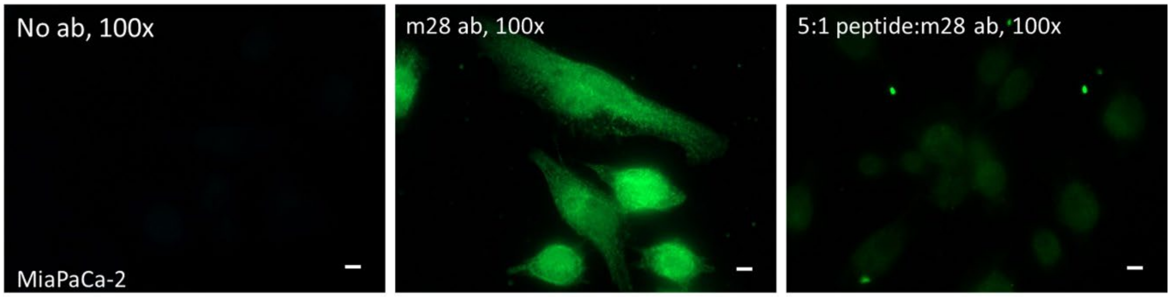


**C** **N-terminal mitochondrial localization signal**

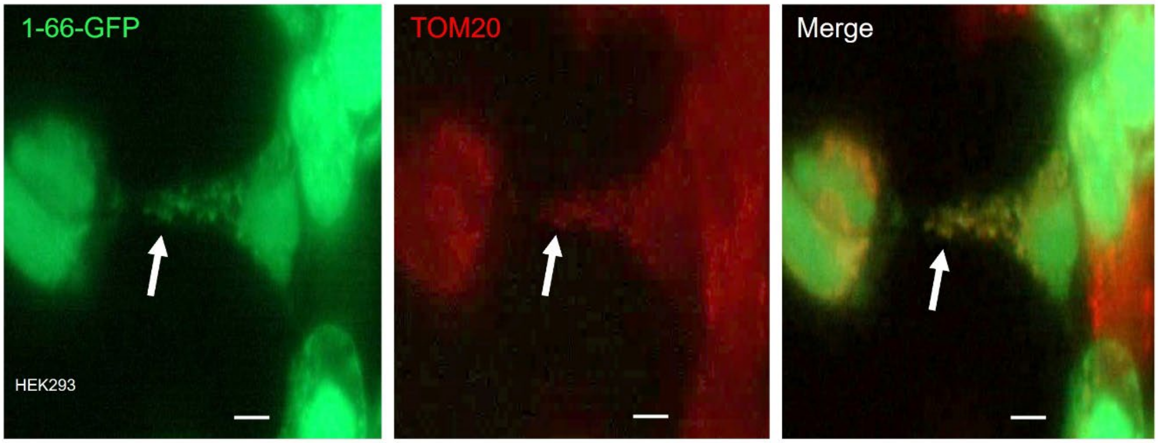

### Figure S3

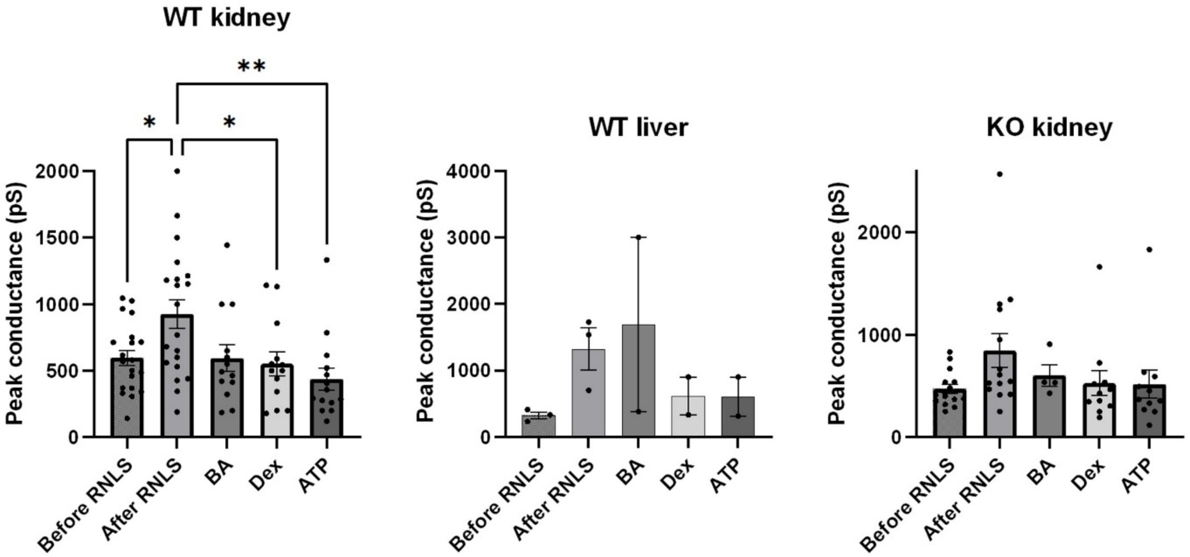
WT kidney
WT liver
KO kidney
Peak conductance (pS)
*
**
0
500
1000
1500
2000
3000
4000
Before RNLS
After RNLS
BA
Dex
ATP

**Figure S4**

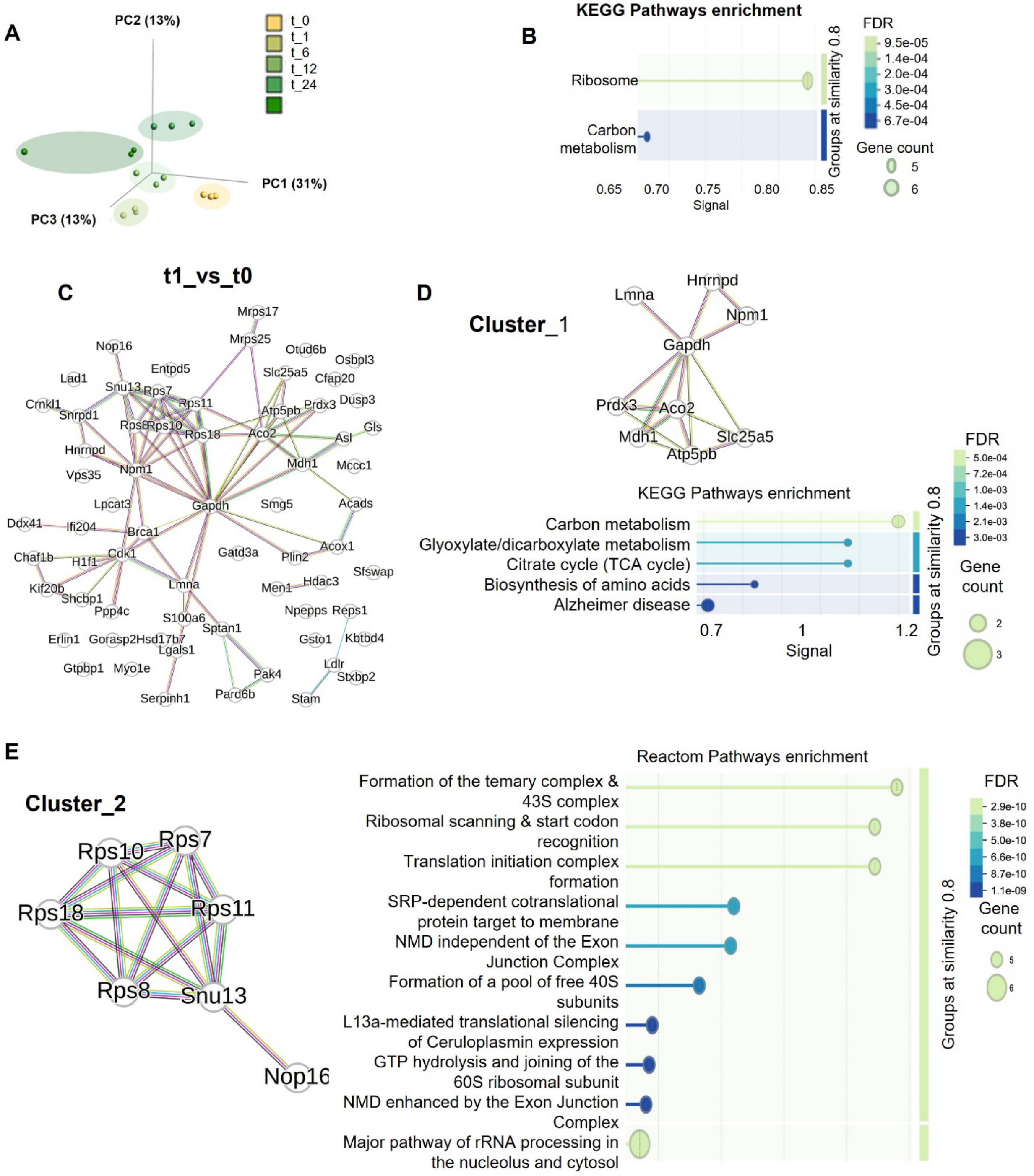
A
PC2 (13%)
PC1 (31%)
PC3 (13%)
t_0
t_1
t_6
t_12
t_24
B
KEGG Pathways enrichment
Ribosome
Carbon metabolism
0.65
0.70
0.75
0.80
0.85
Signal
Groups at similarity 0.8
FDR
9.5e-05
1.4e-04
2.0e-04
3.0e-04
4.5e-04
6.7e-04
Gene count
5
6
C
t1_vs_t0
Mrps17
Mrps25
Nop16
Otud6b
Osbpl3
Entpd5
Slc25a5
Cfap20
Lad1
Snu13
Rps7
Dusp3
Crnkl1
Snrpd1
Rps11
Atp5pb
Prdx3
Rps8
Rps10
Rps18
Aco2
Asl
Gls
Hnrnpd
Mdh1
Mccc1
Vps35
Npm1
Lpcat3
Gapdh
Smg5
Acads
Ddx41
Ifi204
Brca1
Chaf1b
Cdk1
Gatd3a
Acox1
H1f1
Plin2
Sfswap
Kif20b
Shcbp1
Lmna
Men1
Hdac3
Ppp4c
Npepps
Reps1
S100a6
Sptan1
Erlin1
Gorasp2
Hsd17b7
Gsto1
Kbtbd4
Lgals1
Ldlr
Gtpbp1
Myo1e
Pak4
Stxbp2
Serpinh1
Pard6b
Stam
D
Cluster_1
Lmna
Hnrnpd
Npm1
Gapdh
Prdx3
Aco2
Mdh1
Slc25a5
Atp5pb
KEGG Pathways enrichment
Carbon metabolism
Glyoxylate/dicarboxylate metabolism
Citrate cycle (TCA cycle)
Biosynthesis of amino acids
Alzheimer disease
0.7
1
1.2
Signal
Groups at similarity 0.8
FDR
5.0e-04
7.2e-04
1.0e-03
1.4e-03
2.1e-03
3.0e-03
Gene count
2
3
E
Cluster_2
Rps10
Rps7
Rps18
Rps11
Rps8
Snu13
Nop16
Reactom Pathways enrichment
Formation of the temary complex & 43S complex
Ribosomal scanning & start codon recognition
Translation initiation complex formation
SRP-dependent cotranslational protein target to membrane
NMD independent of the Exon Junction Complex
Formation of a pool of free 40S subunits
L13a-mediated translational silencing of Ceruloplasmin expression
GTP hydrolysis and joining of the 60S ribosomal subunit
NMD enhanced by the Exon Junction Complex
Major pathway of rRNA processing in the nucleolus and cytosol
Groups at similarity 0.8
FDR
2.9e-10
3.8e-10
5.0e-10
6.6e-10
8.7e-10
1.1e-09
Gene count
5
6
